# Ultraviolet exposure conditions skin to enhance arbovirus infection and mosquito probing

**DOI:** 10.64898/2026.03.24.713097

**Authors:** Ailish McCafferty-Brown, Amani M Alhazmi, Yonca Keskek Turk, Andrew Durham, Lewis Kajtar, Caoimhe Connolly, Zak Marshall, Daniella A Lefteri, Steven R Bryden, Gerard J Graham, Emilie Pondeville, Kave Shams, Clive S McKimmie

## Abstract

Mosquito-borne viruses are inoculated into skin, yet how environmental exposures shape susceptibility remains unclear. We identify ultraviolet (UV) exposure as an unrecognised determinant of mosquito-borne virus infection. In mice, prior UV exposure increased Semliki Forest virus replication, viraemia and mortality, and also enhanced Zika virus infection. Early susceptibility was driven by recruited CCR2-dependent myeloid cells that were preferentially infected and amplified virus. This phase was transient: by one week, infection shifted toward proliferating fibroblasts within repairing skin, accompanied by a glycolytic tissue signature. In primary human dermal fibroblasts, viral replication was governed by metabolic state rather than proliferation alone, suggesting that repair-associated metabolic reprogramming underlies stromal permissiveness. Topical steroids partially alleviated UV-conditioned enhancement of virus susceptibility. UV also warmed skin and increased mosquito probing. Together, these findings establish UV exposure as a driver of conditioned states that determine arbovirus susceptibility, identifying sunlight as a modifiable determinant of vector-borne disease risk.

**In Brief / eTOC blurb:** McCafferty-Brown et al. show that ultraviolet exposure conditions skin to enhance mosquito-borne virus infection. Susceptibility shifts from infected myeloid cells early after UV to proliferating fibroblasts with a glycolytic tissue signature during repair, while UV-exposed skin also increases *Aedes aegypti* probing.

## Introduction

Mosquito-borne arboviruses infect hundreds of millions of people each year and include dengue virus (DENV), Zika virus (ZIKV) and chikungunya virus (CHIKV), for which the day-biting mosquito *Aedes aegypti* is a major vector^1,2^. Clinical outcome is highly variable, ranging from inapparent infection to severe disease with long-term sequelae. Although viral dose, viral genetics and systemic immune status influence disease outcome, infection is initiated in skin, where virus is deposited into a tissue environment already shaped by prior exposure, inflammation and repair. This local context may influence not only the capacity of virus to replicate after inoculation, but also the cues that determine how mosquitoes interact with the skin surface. The extent to which a pre-existing conditioned state of the inoculation site alters arbovirus susceptibility and vector behaviour remains poorly understood.

The skin inoculation site is not a passive portal of entry. During mosquito feeding, virus and saliva are deposited into the dermis, where salivary factors, tissue injury and viral sensing activate overlapping host pathways^3–5^. These responses can have opposing effects on infection: mosquito biting and bite-induced inflammation can recruit virus-permissive myeloid cells and enhance infection, whereas viral nucleic acid sensing induces type I interferon (IFN) and antiviral restriction^5–8^. Inflammatory recruitment of myeloid cells to the inoculation site has been shown to enhance infection by genetically distinct arboviruses, including flaviviruses, alphaviruses and bunyaviruses^6,9–17^. Because the cellular composition of the inoculation site changes over time, environmental conditioning may influence infection through distinct phases: early inflammatory recruitment of permissive myeloid cells followed by repair programmes dominated by stromal cell remodelling.

Ultraviolet (UV) radiation is one of the most common environmental exposures affecting skin, and arboviruses are often transmitted in regions where human exposure to intense sunlight is high. This overlap is likely to become increasingly relevant as climate warming expands the geographic and seasonal suitability of mosquito vectors and increases outdoor heat and UV exposure^18,19^. UV induces DNA damage, inflammatory signalling, vascular change, immune modulation and tissue repair^20–23^. These responses can persist after the initial exposure has resolved, suggesting that UV may establish a durable conditioned skin state. Sunlight exposure has been linked to altered susceptibility to some skin-associated viral infections, including herpes simplex^21,24,25^, but whether UV exposure alters mosquito-borne virus infection at the exposed inoculation site has not been defined. Nor is it known whether UV-induced changes in local inflammation, perfusion, temperature or skin surface properties influence mosquito probing behaviour.

Dermal fibroblasts are central regulators of the skin response to injury^23,26–28^. Beyond providing structural support, fibroblasts coordinate matrix remodelling, inflammatory signalling, leukocyte recruitment and restoration of tissue architecture during repair. These functions are highly state-dependent: fibroblasts can adopt distinct inflammatory, reparative and matrix-producing programmes according to local cues. This makes them strong candidates for encoding durable tissue memory after environmental damage. Consistent with this concept, recent work has shown that vector-derived inflammatory cues can reprogramme dermal fibroblasts and enhance cutaneous arbovirus infection^29^. Thus, UV-induced fibroblast remodelling may provide a mechanism through which prior environmental exposure alters subsequent viral permissiveness.

Tissue repair is also a metabolic process. Dermal fibroblasts are central executors of repair, coordinating matrix remodelling, proliferation and restoration of tissue architecture. These functions require substantial metabolic adaptation, including shifts in glycolytic and mitochondrial pathways that support biosynthesis and stress resilience^30,31^. Such changes are directly implicated in viral replication, because many viruses depend on host metabolic pathways to generate the energy, nucleotides, lipids and membranes required for replication^32–37^. This may be especially relevant for alphaviruses: Semliki Forest virus (SFV) and Sindbis virus require glycolysis for optimal replication^33^, while SFV and Ross River virus can induce pro-viral metabolic changes through PI3K/AKT signalling^32^. UV exposure can also alter glucose consumption and lactate production in human skin fibroblasts^38^. Together, these observations suggest that UV-induced repair may create a metabolically permissive stromal niche for arbovirus amplification.

If UV-conditioned skin alters the cellular and metabolic state of the inoculation site, it may also alter the cues that determine whether mosquitoes probe that site. *Ae. aegypti* integrates olfactory, visual, humidity and thermal signals during host seeking^39–41^, and thermal cues can enhance host-seeking behaviour when combined with other host-associated signals^42^. Since UV exposure can increase inflammation, blood flow and local tissue temperature, UV-conditioned skin could plausibly influence both sides of vector-borne transmission: the probability that virus amplifies after inoculation, and the likelihood that mosquitoes probe exposed skin.

Here, we show that UV exposure establishes a local conditioned skin state that enhances mosquito-borne virus infection. Using an immunocompetent mouse model incorporating *Ae. aegypti* bites, we find that erythemal and non-erythemal UV exposures increase arbovirus replication and dissemination from the exposed site. Mechanistically, susceptibility evolves over time: early after UV exposure, recruited CCR2-dependent myeloid cells amplify infection, whereas by one-week post-UV exposure, infection shifts toward proliferating fibroblasts within repairing skin and is accompanied by a glycolytic tissue signature. Experiments in primary human dermal fibroblasts support metabolic state, rather than proliferation alone, as a determinant of cellular permissiveness. UV exposure also increases skin temperature and mosquito probing. Together, these findings identify sunlight exposure as a modifiable environmental determinant of arbovirus susceptibility and vector-borne disease risk.

## Results

### UV exposure at the inoculation site enhances arbovirus infection, dissemination and disease

To define whether prior UV exposure alters susceptibility to mosquito-borne virus infection, we used an established immunocompetent mouse model in which arbovirus is inoculated into *Ae. aegypti*-bitten skin^6^. This approach preserves the bite-conditioned tissue environment that shapes early infection, including changes in local leukocyte recruitment and tissue fluid balance^12^, while allowing controlled delivery of virus at the inoculation site. Because many medically important arboviruses replicate poorly in fully immunocompetent mice^5^, we primarily used Semliki Forest virus (SFV) as a tractable model alphavirus ^6,43,44^ with Zika virus (ZIKV) used to test broader relevance^8,12^.

Mice received a single erythemal UVA/B exposure to dorsal foot skin^45^, an accessible site for localised UV exposure and mosquito biting that, unlike most truncal mouse skin, contains epidermal and dermal melanocytes^46–48^. 24 hours later, UV-exposed or unexposed skin was exposed to *Ae. aegypti* bites and inoculated with SFV (Figure 1A, S1A). Prior UV exposure increased viral RNA at the inoculation site, viral RNA in spleen, and infectious virus in blood at 24 h post-infection. Mosquito biting alone enhanced infection, as expected^6^, and UV exposure in the absence of biting also increased viral burden compared with resting skin. However, the combination of prior UV exposure and mosquito biting produced the highest viral loads (Figure 1A, S1A). UV-enhanced susceptibility was UVB-dependent, as UVA alone did not reproduce this effect (Figure S1B). Prior UV exposure also increased ZIKV RNA and infectious virus in blood, indicating that the effect was not restricted to SFV (Figure 1B).

**Figure 1.**
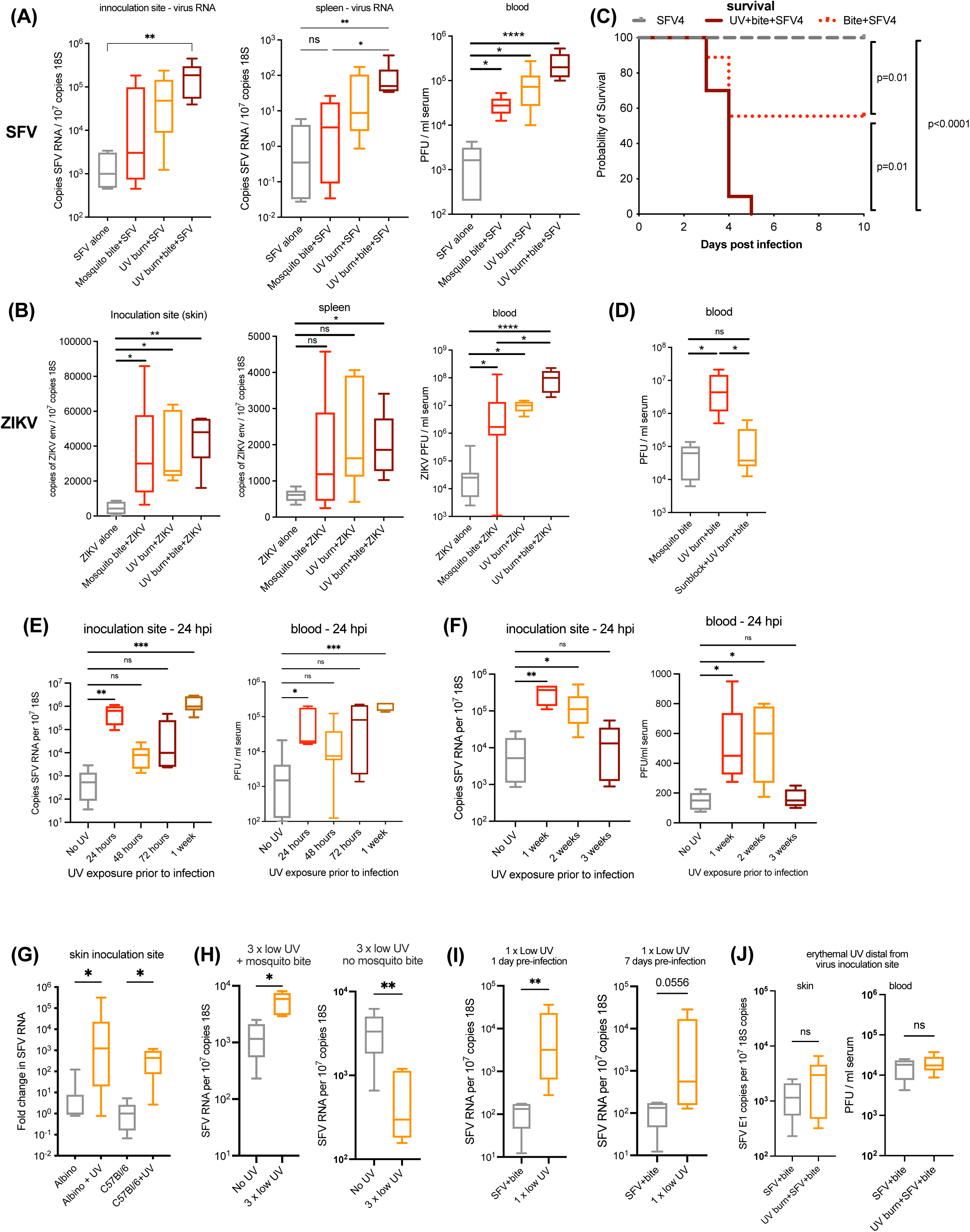
Prior ultraviolet (UV) exposure of the skin enhances host susceptibility to arbovirus infection. (A-B) Mice were exposed to a single erythemal dose of UV radiation (2 MED; 400 mJ/cm²) on the dorsal foot skin. 24 h later, exposed skin was either left unbitten or exposed to *Ae. aegypti* bites and immediately infected with 1×10⁴ PFU SFV (A) or 1×10⁴ PFU ZIKV (n = 6). Viral RNA and infectious virus were determined at 24 h post-infection (hpi). (C) Survival of mice infected with 1×10⁴ PFU SFV at UV-exposed or control skin; mice were monitored until humane endpoints were reached (n = 10). (D) Broad-spectrum SPF50+ sunblock containing UVA and UVB filters was applied topically to the planned UV exposure site 20 minutes prior to erythemal UV exposure. Exposed skin was mosquito bitten and infected with SFV 24 h later (n = 5). (E-F) Mice were infected with SFV at a range of time points post-erythemal UV exposure (n = 6). (G) Syngeneic albino mice and pigmented (C57BL/6) mice were infected with SFV 24 h after an erythemal UV exposure (n = 9). (H) Mice were exposed to repeated non-erythemal UV exposures (0.2 MED; 20 mJ/cm² every 48 h for 6 days) and simultaneously exposed to mosquito bites and infected with SFV 24h after the final exposure (n = 5). (I) Mice were exposed to a single non-erythemal UV exposure and infected with SFV at a mosquito bite 24 h or 7 days later (n = 5). (J) Mice received erythemal UV exposure on one foot and were inoculated with SFV in the unexposed contralateral foot 24 h later (n = 5). All viral RNA in tissues was quantified at 24 hpi by qPCR and normalised to 18S RNA, and infectious virus in serum by plaque assay at 24 hpi. Bars represent the median +/- interquartile range. Statistical significance: \**p*<0.05, \*\**p*<0.01, \*\*\**p*<0.001, \*\*\*\**p*<0.0001, ns = not significant (Mann Whitney or Kruskal–Wallis with Dunn’s post-test for comparison with three or more groups; log-rank Mantel–Cox test for survival analysis).

Enhanced early viral burden translated into worse clinical outcome. SFV infection of UV-exposed skin significantly reduced survival compared with infection of unexposed skin (Figure 1C, S1C). Despite higher early viral loads, serum from UV-exposed mice showed reduced neutralising activity at later time points (Figure S1D), suggesting that increased early replication did not result in improved protective humoral activity.

UV-enhanced infection required local exposure of the inoculation site. SPF50+ sunblock prevented the effect of UV exposure (Figure 1D). The effect was also temporally regulated, with the magnitude of UV-induced susceptibility dependent on the interval between exposure and infection. Infection at 48–72 h post-UV produced non-significant increases in viral load, whereas infection at 1–3 weeks post-UV significantly increased viral burden (Figure 1E,F). Consistent with this extended susceptibility window, mice infected with ZIKV one week after UV exposure rapidly succumbed within 24 h post-infection (Figure S1F). The presence of melanocytes was dispensable in this model, as UV enhanced infection to a similar extent in pigmented C57BL/6 skin and syngeneic albino skin (Figure 1G).

Repeated non-erythemal UV exposure also enhanced infection at mosquito bite sites (Figure 1H, S1H). This regimen induced photoadaptive changes, including increased epidermal thickening (Figure S1G)^45^. Even a single non-erythemal exposure was sufficient to increase susceptibility (Figure 1I). In contrast, low-dose UV in the absence of a mosquito bite was partially protective (Figure S1I), indicating that UV does not simply increase basal viral permissiveness, but also alters how skin responds to subsequent mosquito biting. Finally, UV acted locally rather than systemically, as infection of unexposed distal foot skin was not enhanced by either erythemal or repeated non-erythemal UV exposure at the contralateral site (Figure 1J, S1E).

Together, these data show that UV exposure conditions the skin inoculation site in a manner that enhances local viral amplification, systemic dissemination and clinical disease. This identifies UV exposure as an external environmental factor capable of modulating skin-based host responses to mosquito-borne virus inoculation.

### At 24 h post-UV, enhanced susceptibility depends on the in vivo tissue environment rather than IFN suppression

UV exposure can induce cell death, damage-associated signals^49^, inflammatory and vasoactive mediators^50–53^ and leukocyte recruitment ^54^. We therefore asked whether UV enhanced viral susceptibility through a direct effect on skin cells, or through processes that require the intact in vivo tissue environment. To test this, skin was either exposed to erythemal UV in vivo and biopsied 24 h later for ex vivo infection, or biopsied first, exposed to the same UV dose ex vivo, and infected 24 h later (Figure S2A). In both settings, viral RNA levels were similar between UV-exposed and control biopsies at 24 hpi, indicating that the early UV-enhanced phenotype was not reproduced by isolated tissue infection.

We next asked whether enhanced infection reflected UV-mediated suppression of adaptive immunity or impaired antiviral IFN induction. UV is a potent modulator of T cell responses, but prior UV exposure still enhanced infection in NOD Scid Gamma mice, which lack lymphoid cells (Figure S2B). Thus, the early increase in susceptibility did not require adaptive immune suppression. We then examined type I IFN and IFN-stimulated gene (ISG) expression after mosquito biting or viral infection. Rather than suppressing these responses, both erythemal and repeated low-dose UV exposure primed skin for similar or increased expression of *ifnb1* and prototypic ISGs after mosquito biting (Figures 2A, S2C) or SFV infection (Figures 2B, S2C–G). UV also enhanced SFV infection in *Ifnar1*-deficient mice (Figure S2H), and enhanced ZIKV infection in the *Ifnar1*-deficient model used above (Figure 1B), indicating that UV-induced susceptibility does not depend on intact type I IFN signalling.

**Figure 2.**
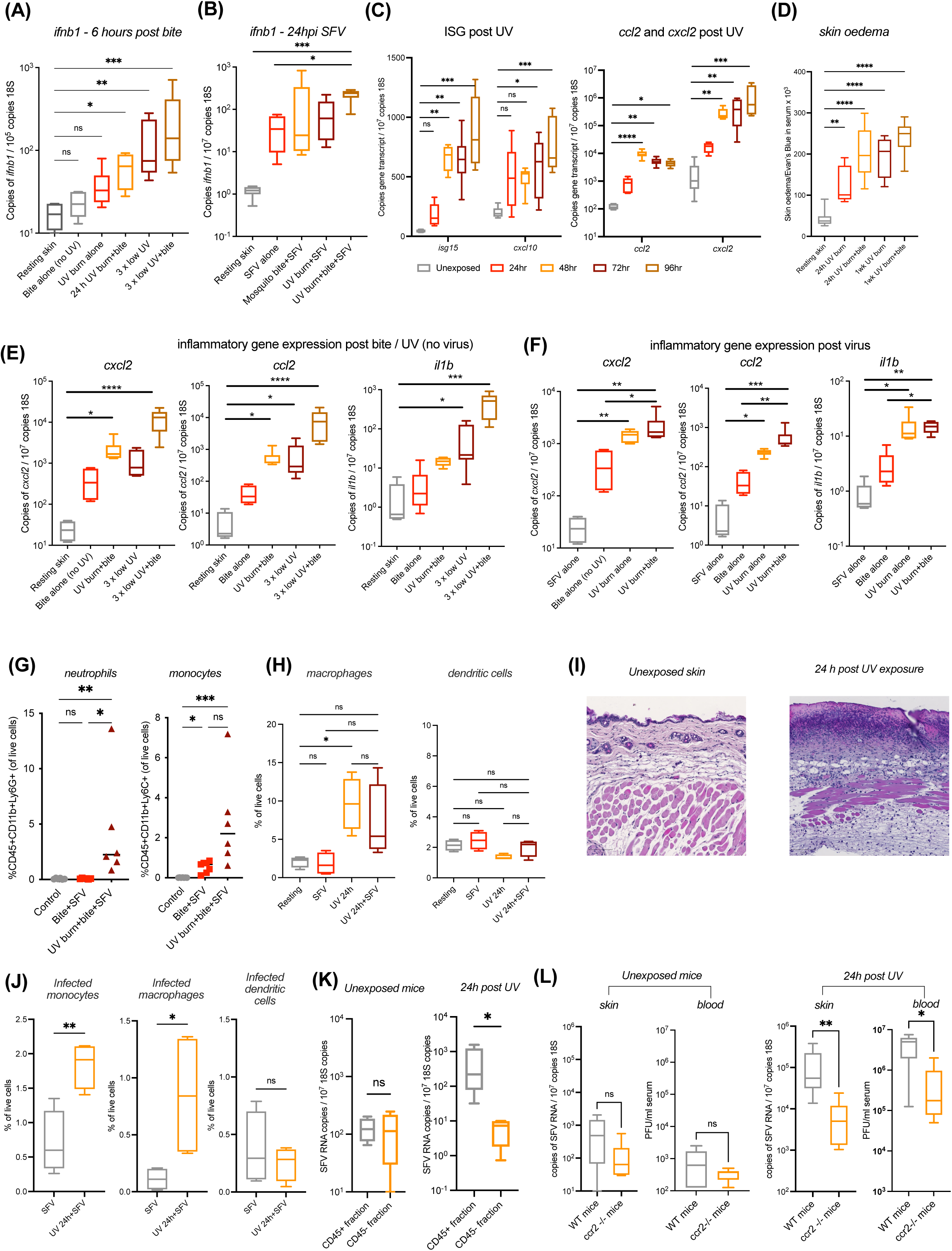
Erythemal UV exposure promotes early susceptibility to arbovirus infection through recruitment and infection of permissive myeloid cells. (A–C) Mice were exposed to either erythemal UV or non-erythemal UV every 48 h for 6 days, and 24 hours after last UV dose, either mosquito bitten alone, or additionally infected with SFV (n = 6). Relative expression of *ifnb1* (A, 6h post bite; B, 24 hpi with bite and SFV) and IFN-stimulated genes (C, 24 hpi with bite and SFV, with times indicating time of infection post-UV exposure), were quantified by qPCR. (D) Erythemal UV exposed mice were administered with Evan’s blue and skin oedema quantified (n = 6). (E,F) Mice were exposed to either erythemal UV or repeated non-erythemal UV and 24 hours after last UV dose, exposed to either mosquito biting alone (E, 6 hours), or additionally infected with SFV4 (F, 24 hpi) and gene expression assessed by qPCR (n = 6). (G–H) Flow-cytometric quantification of leukocytes in skin 24 h after virus infection (n = 6). (I) Mice were exposed to erythemal UV and at 24 h post exposure skin histology was assessed by H&E, with bars representing 100 μm. (J) SFV6-mCherry was inoculated (1×10^3^ PFU) at the exposure site 24 h post-erythemal UV exposure. At 16 hpi, mCherry+ve leukocyte frequencies were assessed by flow cytometry (n = 4). (K,L) Mice (K, WT; L, WT or *ccr2* null) were either exposed to erythemal UV or left unexposed and infected with SFV 24 h later at a mosquito bite. (K) At 6hpi, skin cells were isolated using magnetic separation of CD45 expressing cells and virus RNA quantified by qPCR (*n* = 4). (L) At 24hpi, quantity of virus RNA in tissues were assessed by qPCR and infectious units in blood assessed by plaque assay (n=5, WT; n=6, *ccr2* null). All gene transcripts were quantified by qPCR normalised to 18S RNA. Data represent median ± interquartile range. Groups significantly different from controls or unexposed skin are indicated: Mann Whitney or Kruskal–Wallis with Dunn’s post-test for comparison with three or more groups: \**p*<0.05, \*\**p*<0.01, \*\*\**p*<0.001, \*\*\*\**p*<0.0001, ns = not significant.

Nevertheless, UV-induced IFN/ISG expression may contribute to the temporal pattern of susceptibility. Infection at 48–72 h post-UV produced only modest or non-significant enhancement (Figure 1E), coinciding with elevated *ifnb1* and ISG expression induced by UV alone (Figures 2C, S2I). Thus, UV exposure does not enhance infection by simply suppressing antiviral IFN responses. These data indicate that the early 24 h post-UV susceptibility phenotype is not explained by direct sensitisation of isolated skin, adaptive immune suppression or impaired IFN induction. Instead, it depends on processes present in the intact in vivo inoculation site.

### Early UV-conditioned susceptibility is driven by recruitment and infection of CCR2-dependent myeloid cells

We next asked how erythemal UV exposure enhances susceptibility during the early 24 h post-exposure window. Because UV did not suppress IFN induction, we hypothesised that early susceptibility instead reflected changes in the cellular composition of the inoculation site. In particular, mosquito biting is known to recruit virus-permissive monocytes and macrophages to skin, where they can become infected and amplify arbovirus replication^6,13,14,17^. This recruitment is promoted by bite-induced vascular leakage and oedema^12^.

Erythemal UV exposure similarly induced skin oedema and increased expression of myeloid cell–recruiting chemokines, priming the tissue for an amplified chemokine response to both mosquito biting and viral infection (Figure 2D–F, S3A). Consistent with this, UV exposure increased the frequency of neutrophils, monocytes and macrophages at virus-infected mosquito bite sites, while dendritic cell frequencies were unchanged (Figure 2G,H, S3B). Histological analysis confirmed tissue disruption and a marked leukocyte infiltrate at 24 h post-UV, although skin thickness was not yet increased at this early time point (Figure 2I, S3C).

To determine whether these recruited cells became infected, we inoculated UV-exposed or unexposed skin with SFV6-mCherry and analysed skin single-cell suspensions at 16 hpi. Prior UV exposure significantly increased the frequency of infected mCherry⁺ monocytes and macrophages, whereas infection of dendritic cells, endothelial cells, fibroblasts and epithelial cells was not increased (Figure 2J, S3D). We then separated CD45⁺ leukocytes from CD45⁻ non-leukocytes by magnetic selection. In UV-exposed skin, viral RNA was enriched in the CD45⁺ fraction compared with the CD45⁻ fraction, whereas in unexposed skin viral RNA was distributed similarly between the two compartments (Figure 2K, S3E). Thus, early after UV exposure, increased viral burden was associated specifically with infection of the leukocyte compartment.

We next tested whether UV made these cells intrinsically more permissive, or instead increased the abundance of permissive cells in vivo. CD45⁺ and CD45⁻ cells isolated from UV-exposed or unexposed skin and infected ex vivo showed no UV-dependent increase in susceptibility (Figure S3E). This indicated that the early phenotype was not explained by a cell-autonomous increase in viral permissiveness, but by altered cellular composition of the inoculation site. To test whether CCR2-dependent monocyte recruitment was required, we infected *Ccr2*-null mice, which lack circulating inflammatory monocytes and have reduced monocyte-derived macrophage populations in skin. In contrast to wild-type mice, *Ccr2*-null mice were protected from UV-enhanced infection, with markedly lower viral RNA in skin and infectious virus in blood at 24 hpi after UV exposure. In the absence of UV, *Ccr2*-null and wild-type mice showed similar susceptibility to infection (Figure 2L).

Together, these data show that, at 24 h post-UV exposure, the conditioned inoculation site contains increased numbers of CCR2-dependent myeloid cells that become infected and amplify virus. This provides an early cellular mechanism for UV-enhanced viral replication and dissemination, while leaving open the possibility that later phases of UV-conditioned susceptibility are driven by distinct repair-associated mechanisms.

### By one-week post-UV, myeloid cells become refractory to infection and no longer drive enhanced susceptibility

We next investigated whether the early myeloid-cell mechanism also explained the sustained increase in susceptibility observed when infection occurred one to two weeks after UV exposure (Figure 1E,F). Infection of UV-exposed skin at these later time points still induced elevated *ifnb1* expression (Figure 3A), indicating that enhanced susceptibility was not associated with loss of type I IFN induction. We therefore asked whether persistent recruitment of virus-permissive myeloid cells accounted for the later phenotype.

**Figure 3.**
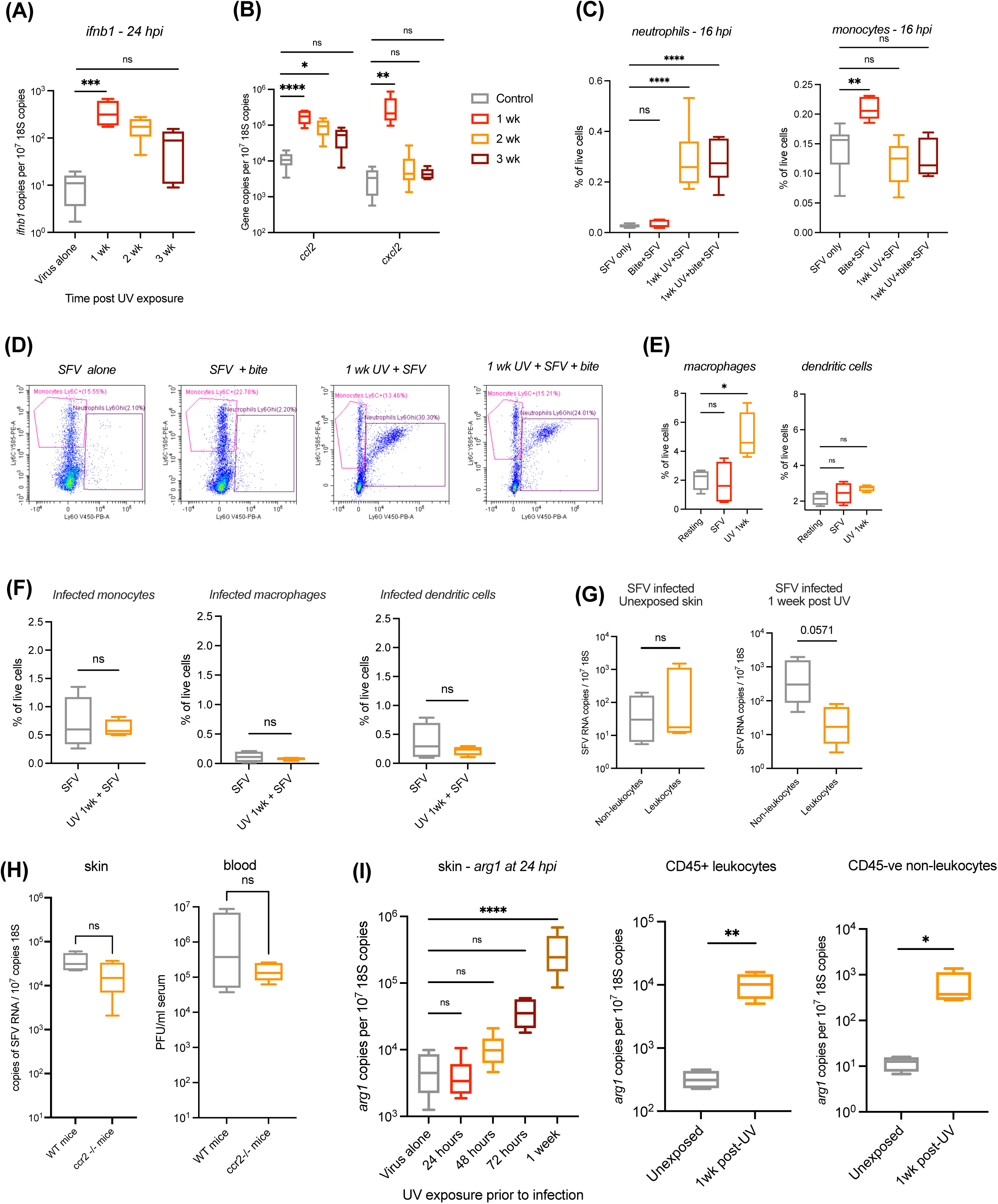
By one week post-UV, myeloid cells become refractory to infection and the tissue shifts toward an Arg1-expressing repair state. (A,B) Expression of *ifnb1* (A) and chemokine (B) transcripts following infection with SFV at 1–3 weeks post-erythemal UV exposure, all sampled at 24 hpi (n = 6). (C–E) Mice were exposed to erythemal UV and 1 week later infected with SFV at mosquito bites. Leukocyte frequencies at 16 hpi were assessed by flow cytometry (n = 6). (F) Mice were exposed to erythemal UV and 1 week later infected with SFV6-mCherry at mosquito bites. At 16 hpi, frequencies of mCherry+ve leukocytes were assessed by flow cytometry (n = 4). (G,H) Mice (G, WT; H, WT or *ccr2* null) were either exposed to erythemal UV or left unexposed and infected with SFV 1 week later at a mosquito bite. (G) At 6hpi, skin cells were separated in to CD45+ (leukocytes) and CD45- (non-leukocytes) using magnetic separation of CD45 expressing cells and virus RNA quantified by qPCR (n = 4). (H) At 24hpi, quantity of virus RNA in tissues were assessed by qPCR and infectious units in blood assessed by plaque assay, showing *ccr2* knockout mice did not exhibit protection from UV-enhanced infection (n=5, WT; n = 6 *ccr2* null). (I) Mice were exposed to erythemal UV and at indicated time post exposure infected with SFV at mosquito bites. *Arg1* transcripts were quantified in either whole tissue biopsies (n = 6) or in skin cell fractions isolated by magnetic labelling of CD45 (n = 4). All gene transcripts were quantified by qPCR normalised to 18S RNA. Bars represent the median +/- interquartile range. Statistical significance: \**p*<0.05, \*\**p*<0.01, \*\*\**p*<0.001, \*\*\*\**p*<0.0001, ns = not significant (Mann Whitney or Kruskal–Wallis with Dunn’s post-test for comparison with three or more groups

Expression of the myeloid cell–recruiting chemokines *ccl2* and *cxcl2* remained elevated at one-week post-UV, although *cxcl2* returned to baseline by week two (Figure 3B). Leukocyte profiling showed increased numbers of neutrophils and macrophages at one-week post-UV, whereas monocyte and dendritic cell frequencies were comparable to those in unexposed, SFV-infected skin (Figure 3C–E). However, unlike at 24 h post-UV, recruited myeloid cells in UV-conditioned skin had become largely refractory to infection: prior UV exposure no longer increased the frequency of infected leukocytes, and the average frequency of infected macrophages was only 0.1% of live cells, more than tenfold lower than in skin infected 24 h after UV exposure (Figure 3F; compare Figure 2J). Consistent with this shift, viral RNA in isolated leukocytes from UV-exposed skin was not elevated and instead showed a trend toward enrichment in the CD45⁻ non-leukocyte fraction (Figure 3G).

Thus, although UV-exposed skin retained increased macrophage numbers (Figure 3E), these cells had become largely refractory to infection and were no longer the dominant infected compartment. In line with this, *Ccr2*-null mice were not protected from UV-enhanced infection when challenged one week after UV exposure (Figure 3H), in contrast to the protection observed at 24 h post-UV (Figure 2L). Together, these data indicate that the later phase of UV-enhanced susceptibility is not dependent on CCR2-dependent myeloid cell amplification.

Instead, infection of UV-conditioned skin one week after exposure was accompanied by marked upregulation of *Arg1* in both CD45⁺ and CD45⁻ cellular fractions (Figure 3I). *Arg1* encodes arginase-1, a marker associated with alternatively activated macrophages57 and reparative fibroblast responses^55,56^. Its induction across leukocyte and non-leukocyte compartments suggests that, by one-week post-UV, the inoculation site has transitioned toward a repair-associated tissue state involving both macrophage polarisation and stromal activation.

### UV-conditioned repair redirects infection toward proliferating fibroblasts

Since viral RNA showed a trend toward enrichment in the CD45⁻ non-leukocyte fraction of UV-exposed skin at one-week post-UV (Figure 3G), we next examined which non-leukocyte populations were infected at this later time point. Infection frequencies in endothelial and epithelial cells were not increased by prior UV exposure. In contrast, the frequency of infected fibroblasts was significantly increased when mice were infected one week after UV exposure (Figure 4A). Thus, after myeloid cells had become largely refractory to infection, UV-conditioned susceptibility was associated with a shift in viral tropism toward the fibroblast compartment.

**Figure 4.**
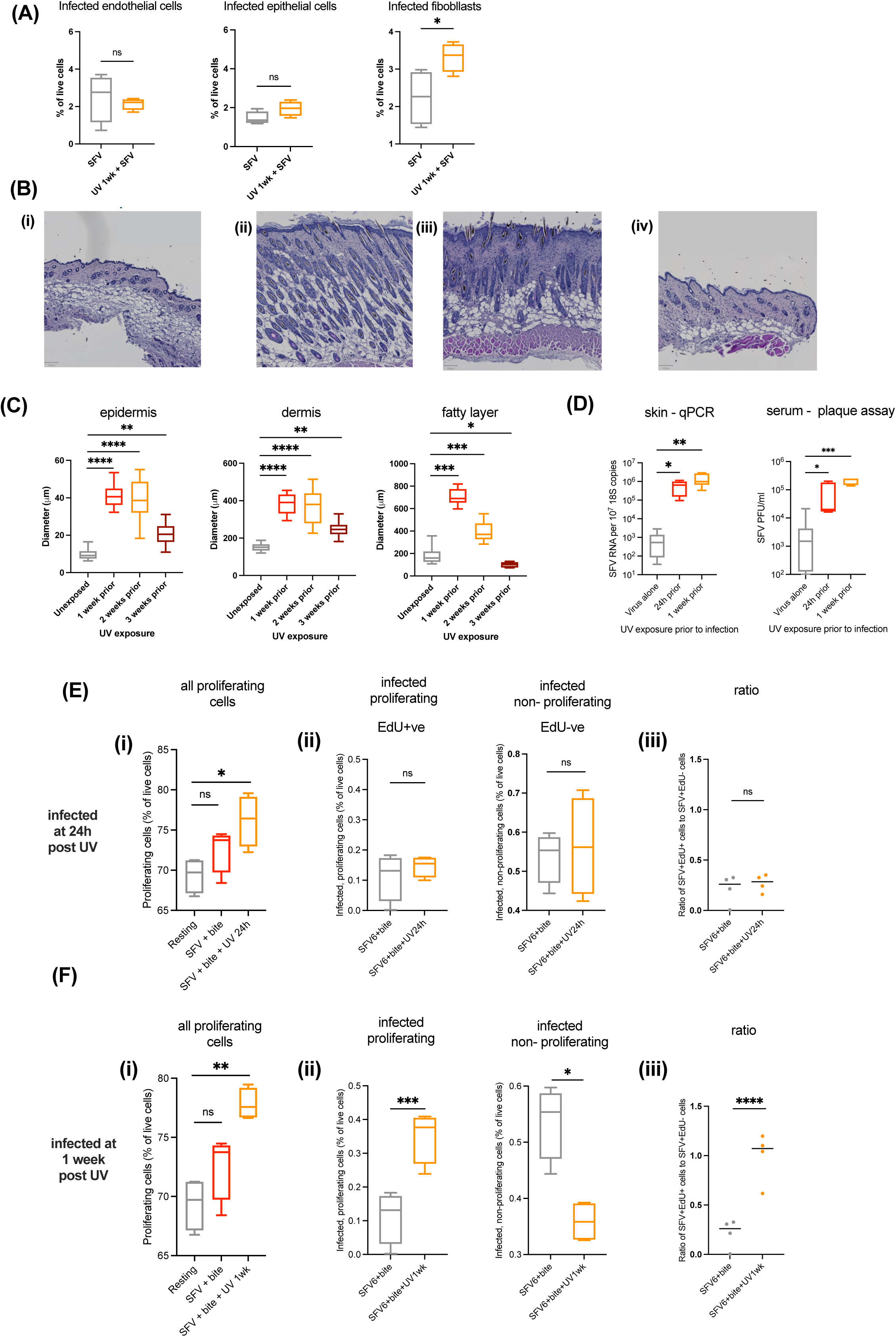
UV-conditioned repair redirects infection toward proliferating fibroblasts. (A-C) Mice were exposed to erythemal UV and at 1 week post exposure and either; (A) infected with SFV6-mCherry and at 24 hpi cells analysed by flow cytometer to define frequency of mCherry^+ve^ endothelial cells, fibroblasts and epithelial cells (n = 4); or (B,C) left uninfected and skin histology assessed by H&E and (C) thickness of each skin compartment defined. Bars representing 100 μm. (D) Mice were exposed to erythemal UV and then at the time points post exposure indicated, infected with SFV6-mCherry (n = 6). (E,F) mCherry-expressing SFV6 and Click-iT EdU labelling were used to identify proliferating (EdU⁺) and infected (mCherry⁺) cells by flow cytometry (n = 4). EdU incorporation and SFV6-mCherry infection frequencies at 24 h post-UV (E) and 1 week post-UV (F). (Ei, Fi) Total numbers of proliferating (EdU^+ve^) cells. (Eii, Fii) Frequencies of double positive mCherry^+ve^EdU^+ve^ cells. (Eiii, Fiii) Ratio of infected proliferating cells to non-proliferating infected cells. Bars represent the median +/- interquartile range. Statistical significance: \**p*<0.05, \*\**p*<0.01, \*\*\**p*<0.001, \*\*\*\**p*<0.0001, ns = not significant (Mann Whitney or Kruskal–Wallis with Dunn’s post-test for comparison with three or more groups).

To define the tissue state associated with this shift, we examined histological sections of UV-exposed skin one week after exposure. UV induced marked thickening of the epidermis, dermis and subcutaneous fat layer (Figure 4B,C), consistent with a broad reparative and hyperplastic response. This was relevant because some viruses, including alphaviruses, can replicate more efficiently in less differentiated or proliferative cellular states than in differentiated non-proliferating cells^57–59^.

We therefore combined SFV6-mCherry infection with EdU labelling to assess whether infection was associated with proliferating cells. Skin from UV-exposed or unexposed sites was infected either 24 h or one week after UV exposure, collected at 24 hpi, digested to single-cell suspensions, and incubated with EdU to identify cells undergoing DNA synthesis. This allowed proliferating EdU⁺ cells and infected SFV6-mCherry⁺ cells to be quantified by flow cytometry. We first confirmed that the SFV6-mCherry infection model reproduced UV-enhanced infection, with viral RNA most strongly elevated when infection occurred one week after UV exposure compared with either unexposed skin or skin infected 24 h after UV exposure (Figure 4D).

This later increase in viral burden was accompanied by an increased number of EdU⁺ cells in UV-exposed skin (Figure 4Ei, 4Fi). When infection occurred 24 h after UV exposure, infected cells showed no clear bias toward the EdU⁺ compartment (Figure 4Eii, 4Eiii). In contrast, when infection occurred one week after UV exposure, virus was preferentially detected in EdU⁺ cells, while infection of non-proliferating cells was reduced (Figure 4Fii, 4Fiii). Thus, the later phase of UV-conditioned susceptibility was characterised by increased infection of fibroblasts and preferential infection of proliferating cells within repairing skin.

Together, these findings show that by one-week post-UV, infection is redirected away from the early myeloid compartment and toward proliferating fibroblasts within a reparative tissue environment. This identifies proliferating fibroblasts as the dominant in vivo correlate of the later UV-conditioned susceptibility state, while leaving open whether proliferation itself, or the metabolic programme associated with repair, drives enhanced permissiveness.

### Fibroblast metabolic state, rather than proliferation alone, determines viral permissiveness

The enhanced infection of proliferating fibroblasts observed in vivo could reflect either intrinsic properties of cycling cells or the altered metabolic state associated with tissue repair. To distinguish between these possibilities, we cultured primary human dermal fibroblasts (HDFs) under conditions that modulated cell cycle progression. Treatment with the cyclin-dependent kinase inhibitor roscovitine reduced *KI67* expression, consistent with reduced proliferation, yet increased SFV RNA levels in a dose-dependent manner at 24 h post-infection (Figure 5A). Similarly, palbociclib, which blocks CDK4/6-mediated G1/S transition, also increased viral RNA accumulation (Figure 5B). Low-serum culture conditions, which reduced Ki67 staining compared with high-serum controls (Figure S4), markedly enhanced viral susceptibility, producing multi-log increases in SFV RNA (Figure 5C). Likewise, high-confluency cultures, typically characterised by contact inhibition and reduced proliferation, supported greater viral replication than medium-confluency cultures (Figure 5D). Together, these findings indicate that proliferation status alone does not determine fibroblast permissiveness to virus infection.

**Figure 5.**
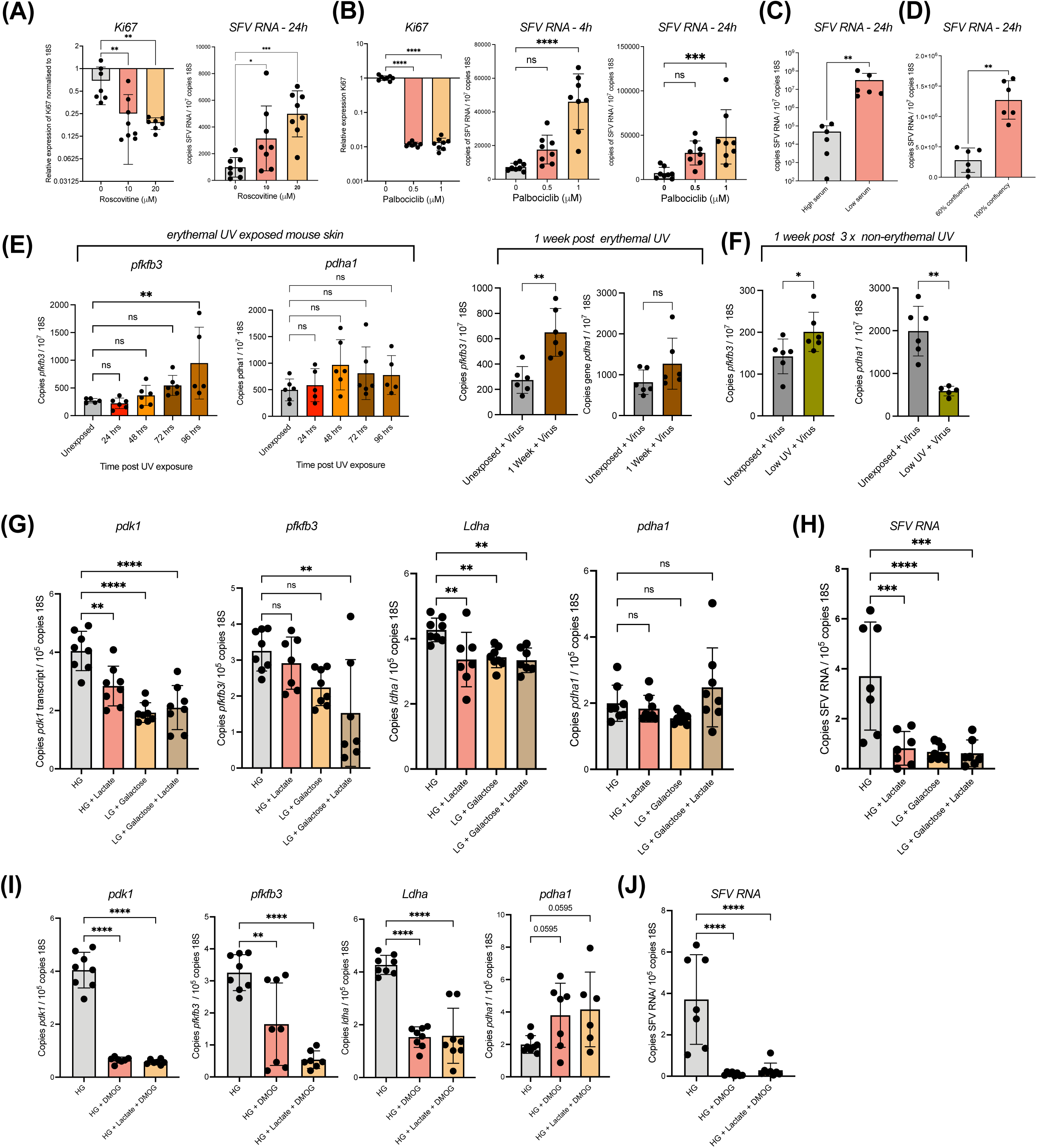
Fibroblast metabolic state, rather than proliferation alone, determines viral permissiveness. (A,B) Primary human dermal fibroblasts (HDFs) were treated with increasing concentrations of roscovitine (A) or palbociclib (B) to inhibit cell cycle progression, then infected with SFV (MOI = 2 for 4 hpi; MOI 0.05 for 24 hpi). Ki67 expression was assessed by qPCR in uninfected controls (n = 8). (C) HDFs were cultured in high serum (10% FBS) or low serum (0.5% FBS) conditions for 24 h before SFV infection (MOI=0.5, 24 hpi, n = 8). (D) HDFs were seeded at either high confluency and left for 5 days to establish cell cycle contact inhibition, or seeded at lower confluency, and infected with SFV (MOI=0.5) and virus RNA quantified at 24 hpi (n = 6). (E,F) Mice were exposed to either single erythemal UV (E), or repeated non-erythemal UV (F), and metabolic gene expression assessed at indicated time points (n = 6). (G,H) HDFs were cultured in standard high-glucose (HG) media or low-glucose (LG) media with or without D-galactose (10 mM) and L-lactate (10 mM) for 48 h prior to SFV infection. (G) Metabolic gene expression and (H) virus RNA levels at 24 hpi (n = 8). (I,J) HDFs were treated with DMOG (0.5 mM) alone or with lactate (10 mM) for 24 h before SFV infection. (I) Glycolytic genes and PDHA1 and (J) Virus RNA quantities were defined at 24 hpi (n = 8). All gene transcripts were quantified by qPCR normalised to 18S RNA. Each dot represents a biological sample. Bars represent mean ± SD and statistical comparisons used ordinary one-way ANOVA or unpaired t-test with Welch’s correction. For non-normally distributed viral RNA data, bars represent median ± interquartile range and statistical comparisons used Mann-Whitney test. Statistical significance: *p<0.05, **p<0.01, ***p<0.001, ****p<0.0001, ns = not significant.

We next asked whether metabolic reprogramming associated with UV-induced tissue repair could provide an alternative explanation for the increased infection of proliferating fibroblasts in vivo. Analysis of key metabolic genes in UV-exposed mouse skin revealed significant upregulation of *pfkfb3*, encoding 6-phosphofructo-2-kinase/fructose-2,6-bisphosphatase 3, a regulator of glycolytic flux^60^, by 96 h post-erythemal UV exposure in uninfected skin (Figure 5E). *Pfkfb3* expression was further elevated at one week post-UV in virus-infected skin, coinciding with peak susceptibility. This was not simply due to a generalised increase in metabolic gene expression, as *pdha1*, encoding the pyruvate dehydrogenase alpha subunit that channels pyruvate into mitochondrial oxidative metabolism, remained unchanged (Figure 5E). Repeated non-erythemal UV exposure also increased *pfkfb3* expression by one week, while *pdha1* was decreased (Figure 5F), consistent with wound-associated remodelling of skin metabolic programmes^61,63,65^.

To test whether dermal fibroblast metabolic state directly modulates viral susceptibility, we cultured primary HDFs under conditions designed to reduce the glycolytic gene programme^62^. Lactate supplementation of high-glucose media, provision of galactose as an alternative carbon source, and culture in low-glucose conditions reduced expression of the glycolytic genes *pdk1*, *ldha* and, in galactose plus lactate conditions, *pfkfb3*, while *pdha1* expression remained largely stable (Figure 5G). Critically, conditions that suppressed glycolytic gene expression also reduced SFV replication, with lactate supplementation and low-glucose conditions yielding significantly lower viral RNA than standard high-glucose media (Figure 5H).

To further test whether altering dermal fibroblast metabolism affects viral susceptibility, we treated HDFs with dimethyloxalylglycine (DMOG), a small-molecule metabolic modulator. DMOG, either alone or combined with lactate, produced the strongest suppression of the measured glycolytic gene programme, with multi-fold decreases in *pdk1*, *ldha* and *pfkfb3*, alongside a trend toward increased *pdha1* expression (Figure 5I). Correspondingly, DMOG-treated cells showed the greatest resistance to viral infection, with multi-fold reductions in viral RNA (Figure 5J).

Together, these data indicate that fibroblast permissiveness is governed by metabolic state rather than proliferation alone*. In vivo*, proliferating fibroblasts may therefore mark a repair-associated state in which altered metabolism creates a permissive stromal environment for viral replication.

### Topical corticosteroids partially reduce UV-enhanced dissemination and neuroinflammation

We next asked whether treatment of UV-damaged skin could modify the enhanced susceptibility state. Vitamin D treatment produced modest histological improvement but did not reduce UV-induced neutrophil or monocyte accumulation, skin viral RNA, or serum infectious virus, and was therefore not pursued further (Figure S5A–C). We therefore tested topical clobetasol propionate, a high-potency corticosteroid used in UK dermatology practice for severe sunburn, applied after erythemal UV exposure and stopped one day before infection at the one-week peak of UV-enhanced susceptibility. Topical steroid treatment partially normalised UV-induced changes in skin architecture, restoring epidermal and fatty layer thickness toward baseline (Figure 6A). Steroid-treated UV-exposed mice no longer showed significantly increased skin viral RNA or viraemia at 24 hpi compared with unexposed controls, indicating partial protection from UV-enhanced early dissemination (Figure 6B). By 4 dpi, steroid-treated mice showed reduced viral RNA in skin and spleen, while brain viral RNA was no longer significantly elevated relative to mice infected without prior UV exposure, although values remained variable (Figure 6C). UV exposure also increased perivascular inflammatory infiltrates in the brain, consistent with SFV-associated encephalitic pathology, and this was reduced by topical steroid treatment (Figure 6D,E; Figure S6A). These effects were not explained by reduced neutrophil or monocyte accumulation in skin, nor by increased *Ifnb1* or *Rsad2* expression after infection (Figure S6B,C). Thus, topical corticosteroids partially reduced the downstream consequences of UV-conditioned susceptibility, including early viraemia and neuroinflammation, but did not fully reverse the UV-conditioned infection state.

**Figure 6.**
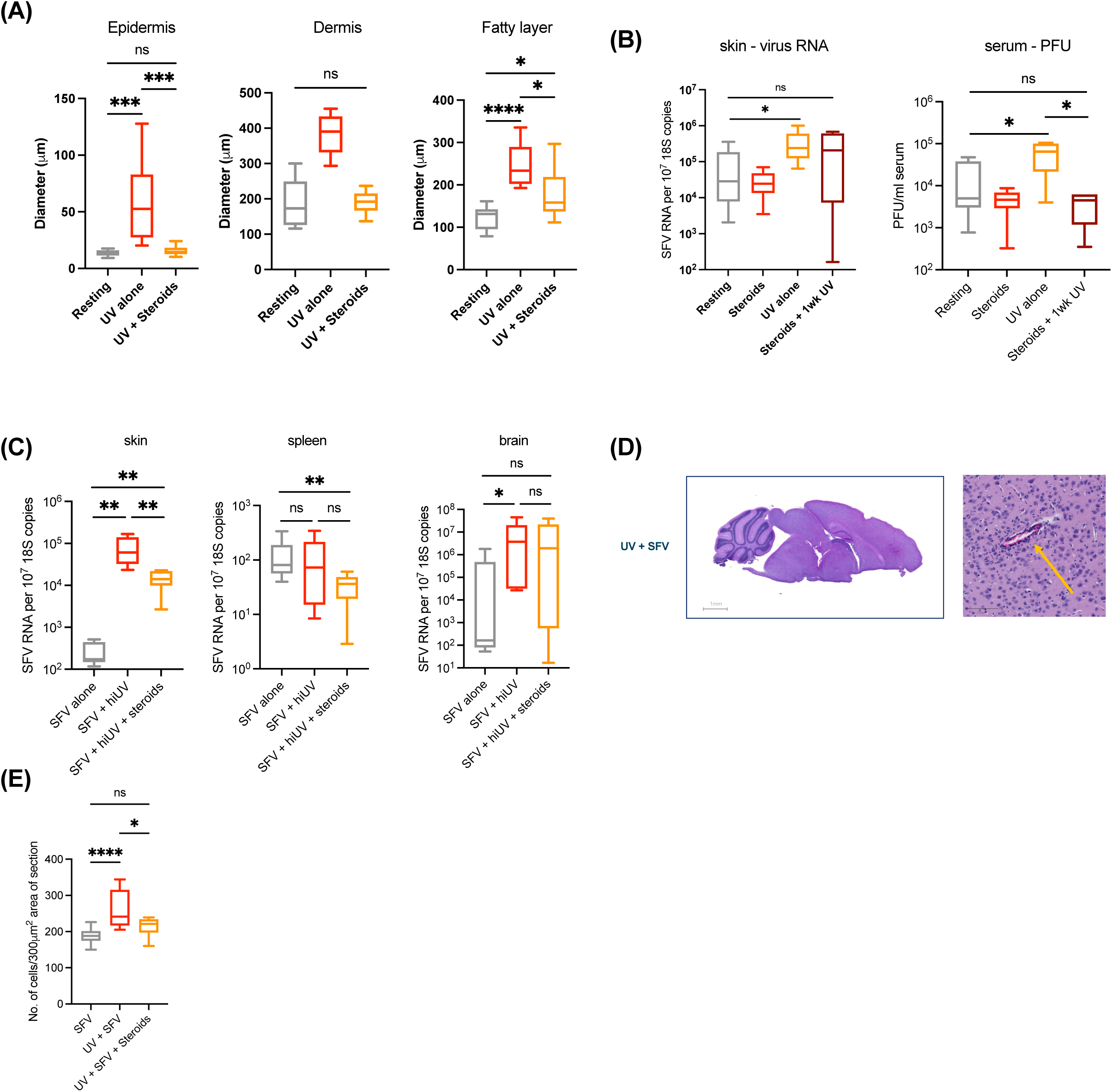
Topical corticosteroids modulate UV-enhanced infection and downstream inflammation. Mice were exposed to erythemal UV and either left untreated or administered with topical clobetasol propionate twice daily. (A) At 1 week post-UV, skin sections were assessed by H&E histology to define thickness of each compartment. (B-E) At 1 week post-UV, mice were infected with SFV at mosquito bites and quantities of virus RNA and blood infectious units defined at either 24hpi (B) or virus RNA at 96 hpi (C) (n = 6). (D) Histology of representative perivascular cuff in the brain cortex. (E) Mono-nuclear infiltrate cells around blood vessels were counted per 300 μm² area of section using QuPATH, taken from 10 areas per mouse, from 3 mice per condition. All viral RNA in tissues were quantified by qPCR and normalised to 18S RNA, and infectious virus in serum by plaque assay. Bars represent the median +/- interquartile range. Statistical significance: \**p*<0.05, \*\**p*<0.01, \*\*\**p*<0.001, \*\*\*\**p*<0.0001, ns = not significant (Mann Whitney or Kruskal–Wallis with Dunn’s post-test for comparison with three or more groups).

### UV exposure increases skin temperature and mosquito probing

Finally, we asked whether UV exposure alters skin properties that influence mosquito behaviour. Mosquitoes use thermal, olfactory and other sensory cues to identify suitable feeding sites^39–42^, and UV-induced changes in local inflammation, perfusion or surface chemistry could therefore affect how mosquitoes interact with exposed skin. To determine whether erythemal UV exposure altered local temperature, we measured UV-exposed mouse skin at 24 h intervals after exposure. Skin temperature increased by approximately 1.5°C at 24 h post-UV, before gradually returning to baseline between 72 and 96 h (Figure 7A). We then assessed mosquito behaviour at the time of peak temperature increase. *Ae. aegypti* mosquitoes were allowed to feed on UV-exposed or unexposed skin 24 h after erythemal UV exposure. Time to first landing was unchanged, indicating that UV exposure did not detectably alter initial landing behaviour. However, both the number of mosquitoes that probed UV-exposed skin and the duration of probing were significantly increased compared with unexposed controls (Figure 7B).

**Figure 7.**
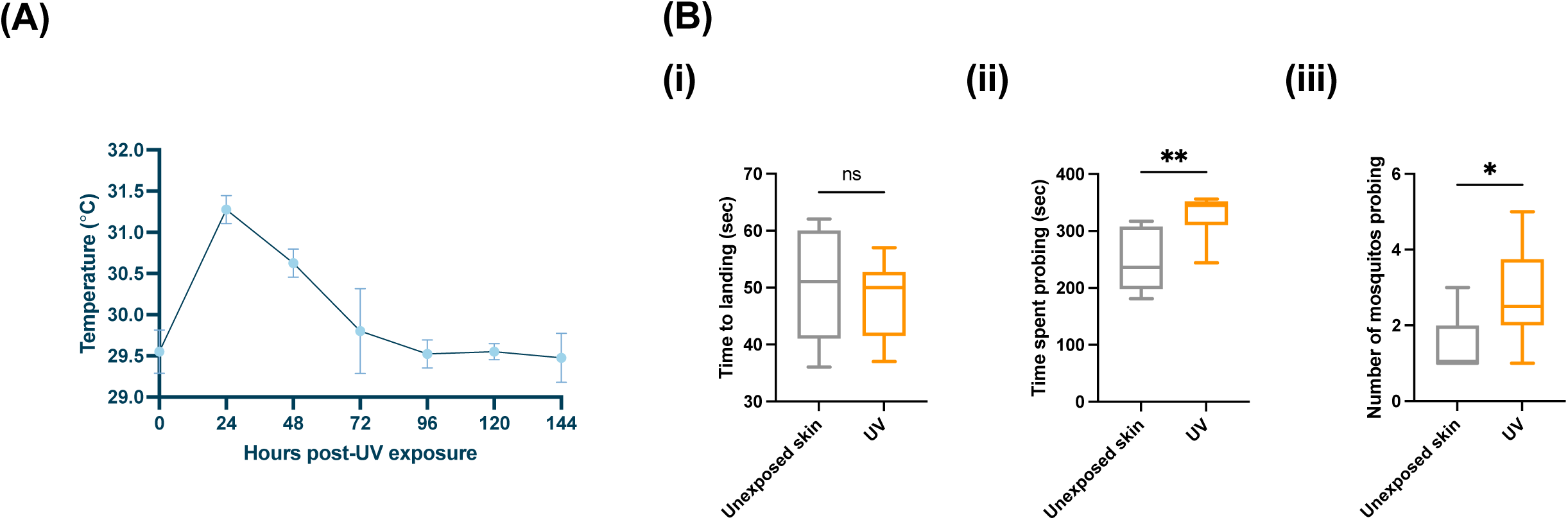
UV exposure increases skin temperature and mosquito probing behaviour. (A) Mice were exposed to erythemal UV and skin temperature at the exposed site measured every 24 h (n = 8). (B) Mice were exposed to erythemal UV. At 24h post-UV, mice were exposed to *Ae. aegypti* mosquitoes at the burn site, with all other body surfaces covered by an insulating impenetrable material. (i) Time from when mouse skin was exposed to cage, to when first mosquito landed on the skin. (ii) Time from when mosquitos began probing to when they were engorged with blood was measured. (iii) Number of mosquitos probing each mouse were counted (n = 8 mice). Bars represent the median +/- interquartile range. Statistical significance: *p<0.05, **p<0.01, ns = not significant (Mann Whitney).

These findings show that erythemal UV exposure alters the local skin environment in a way that increases *Ae. aegypti* probing. This effect may reflect increased skin temperature, UV-induced changes in volatile cues, altered perfusion, or changes in skin surface properties. Thus, UV-conditioned skin may influence not only host susceptibility after virus inoculation, but also mosquito behaviour at the exposed site.

## Discussion

Our findings identify UV exposure as a previously unrecognised environmental determinant of mosquito-borne virus infection. UV conditioned the skin inoculation site to enhance local viral amplification, systemic dissemination and disease, without impairing type I IFN induction or requiring adaptive immune suppression. Instead, UV created a temporally evolving tissue state in which susceptibility changed over time. At 24 h post-UV, recruited CCR2-dependent myeloid cells amplified infection. By one week, these cells had become largely refractory to infection, and susceptibility shifted toward proliferating fibroblasts within repairing skin. Thus, recent UV exposure can shape arbovirus outcome by altering the tissue into which virus is deposited.

Rather than reflecting a single inflammatory mechanism, UV-conditioned susceptibility evolved through distinct cellular phases. At 24 h post-UV, monocytes and macrophages were increased and preferentially infected, and loss of CCR2-dependent recruitment largely protected against UV-enhanced infection. This parallels mosquito biting in unexposed skin, where vascular leakage and inflammatory recruitment increase the frequency of virus-permissive myeloid cells^6,12–15^. However, at one week post-UV, macrophages remained more frequent but were no longer preferentially infected, viral RNA shifted toward the non-leukocyte compartment, and *Ccr2*-deficient mice were no longer protected. UV-conditioned susceptibility is therefore better understood as a sequence of tissue states, rather than a single inflammatory mechanism.

The later phase of this sequence coincided with tissue repair and a shift toward fibroblast infection. UV-exposed skin showed thickening of epidermal, dermal and subcutaneous compartments, increased infection of fibroblasts, and preferential infection of proliferating cells. However, HDF experiments showed that proliferation alone was insufficient to explain permissiveness: pharmacological or culture-based suppression of proliferation did not reduce infection and often increased viral RNA. Instead, viral replication tracked with fibroblast metabolic state. In UV-exposed mouse skin, susceptibility coincided with a glycolytic tissue signature, while in HDFs, suppression of the glycolytic gene programme reduced SFV replication. These data support a model in which repair-associated fibroblast states create a metabolically permissive stromal environment for viral amplification.

This model is further supported by several findings that were unexpected. First, UV-enhanced susceptibility was not linear over time: infection was only modestly enhanced at 48–72 h post-UV, coinciding with UV-induced IFN/ISG expression, before susceptibility increased again during the repair phase. Second, repeated low-dose UV exposure was partially protective in the absence of mosquito biting, indicating that UV does not simply increase basal permissiveness but instead alters how skin responds to vector-associated inoculation. Third, DMOG unexpectedly suppressed, rather than induced, the measured glycolytic gene programme in HDFs and strongly reduced SFV replication. This suggests that fibroblast permissiveness is not explained by HIF activation alone, but by a context-dependent metabolic state that may differ between fibroblasts, immune cells and transformed cell systems.

These findings place fibroblasts as active determinants of arbovirus susceptibility, rather than passive structural targets. Dermal fibroblasts organise tissue repair, inflammatory signalling and matrix remodelling^23,26–28^, and vector-derived inflammatory cues can reprogramme dermal fibroblasts to enhance Toscana virus infection^29^. More broadly, environmental or inflammatory exposures that alter fibroblast state may generate permissive niches for cutaneous arbovirus replication, helping to explain variation in infection outcome and clinical severity.

Topical corticosteroid treatment partially modified the UV-conditioned phenotype. Clobetasol reduced early viraemia, viral RNA in peripheral tissues and perivascular inflammatory infiltrates in the brain, but did not fully restore antiviral resistance or eliminate the effects of prior UV exposure. Thus, suppressing post-UV inflammation reduced some downstream consequences of UV conditioning, but may not completely reverse the repair-associated state that supports susceptibility. This suggests that treatment of sunburn may not fully mitigate the increased susceptibility associated with recent intense UV exposure, although this requires direct testing in human skin.

The consequences of UV conditioning were not restricted to host susceptibility after inoculation. Erythemal UV also increased local skin temperature and altered mosquito feeding behaviour, increasing both the number of *Ae. aegypti* mosquitoes probing the exposed site and the duration of probing, while time to first landing was unchanged. Thus, UV exposure could modify arboviral risk at two levels: by increasing mosquito probing and opportunities for infectious inoculation, and by conditioning the inoculation site to support greater viral amplification, dissemination and disease after virus delivery.

This dual effect has translational implications for prevention, risk assessment and intervention. Sun protection is already promoted to reduce UV-mediated skin damage and skin cancer risk, and our data suggest that UV avoidance may have additional value where mosquitoes transmit arboviruses. Recent UV exposure is unlikely to predict severe disease alone, but could be considered alongside established clinical and laboratory warning signs in outbreak settings where early recognition of deterioration is critical, as in dengue^64^. The findings also support the earliest skin stage of arbovirus infection as a tractable point for intervention. Salivary antigen-based strategies such as AGS-v PLUS show that the human skin response to vector exposure can be reprogrammed^66^; our findings extend this principle by showing that environmental tissue states also determine permissiveness. Interventions targeting skin repair, stromal metabolism or vector-induced tissue responses could therefore modify disease trajectory by targeting the host tissue state that permits early amplification, rather than the virus alone.

Several points will be important for defining how these findings translate to human exposure and transmission. Mouse UV models do not fully capture the heterogeneity of human sun exposure, skin pigmentation, anatomy or repair responses. The metabolic data also have limits: the in vivo signature was measured in whole skin, while mechanistic perturbation was performed in HDFs. Finally, increased probing provides a plausible route to increased inoculation opportunities, but field studies are needed to determine whether this alters transmission risk in humans.

In summary, UV exposure conditions the skin inoculation site to enhance arbovirus infection, dissemination and disease. Susceptibility evolves from an early CCR2-dependent myeloid amplification phase to a later repair-associated fibroblast phase linked to a glycolytic tissue signature and fibroblast metabolic permissiveness. By connecting sunlight exposure to host tissue state, viral replication and mosquito probing, our study identifies UV exposure as a modifiable environmental determinant of vector-borne disease risk and highlights the skin inoculation site as a target for future antiviral, vaccine and public-health strategies.

## Acknowledgments

We gratefully acknowledge the Leeds Biomedical Services and York Biological Services for technical assistance with mouse procedures, and the Leeds Faculty of Medicine & Health Flow Cytometry Facility for their support. CSM and AM-B were funded by the MRC Medical Research Council [MR/N013840/1] and a University of York startup grant. EP was funded by the MRC (MC_UU_00034/8). We thank Dr Kim Robinson (Hull York Medical School, University of York) for the supply of HDFs. We thank Kosar Babani and Abd Abou Swid (Hull York Medical School) for assistance with experimental work during training in RNA extraction and qPCR.

## STAR Methods and materials

### Cell culture, viruses and mice

Baby hamster kidney-21 (BHK-21) cells and C6/36 mosquito cells were grown and SFV4, SFV6-mCherry and ZIKV prepared, as previously described^6^. The pCMV-SFV4 and pCMV-SFV6-mCherry backbone for production of SFV has been previously described^44,67^. Plasmids were electroporated into BHK cells to generate infectious virus. ZIKV was derived from a Recife isolate (ZIKV PE243), kindly supplied by Prof Alain Kohl (MRC-University of Glasgow Centre for Virus research). Human dermal fibroblasts (HDF) were a kind gift from Dr. Kim Robsinson^68^ (Skin Research Centre, Hull York Medical School, University of York, UK). Unless otherwise specified HDFs were cultured in Dulbecco’s Modified Eagle Medium (DMEM, high glucose with sodium pyruvate) at 10% FCS with penicillin/streptomycin and GlutaMAX supplement.

Viruses were grown once in BHK-21 cells, then passaged once in C6/36 cells. Virus stocks were titred and viraemia was quantified using plaque assays. Unless otherwise specified, all mice were 6-8-week-old wild type mice (C57bl/6J). All mice were derived from a locally bred-colony maintained in a pathogen-free facility, in filter-topped cages and maintained in accordance with local and governmental regulations. *Ifnar-/-* mice, bred on a C57BL/6 background, were originally purchased from Jackson Laboratories, USA and then maintained locally. *Ccr2-/-* mice, bred on a C57BL/6 background, were kindly supplied by Dr Marieke Pingen (University of Glasgow).

### Exposure of mouse foot skin to UV radiation

Mouse skin on the upper (dorsal) side of the foot is an accessible, relatively hair-poor site for localised UV exposure and mosquito biting that contains epidermal and dermal melanocytes, unlike e.g. most truncal mouse skin. As such, foot skin better mimics human skin which has similar melanocyte distribution^46–48^, and does not require hair removal prior to UV exposure. For histology experiments, mouse trunk was also used, in which case hair removal was undertaken by shaving and allowed to rest for several days prior to experiments. In all cases, skin was exposed to UV radiation containing both UVA and UVB to more accurately model sun exposure. A 8W UVM-28 mid-range lamp (302 nm peak at 1mW/cm^2^, Ultraviolet Products) was used, which emits most of its energy within the UVB range, with the remaining 20–30% output as UVA. We and others^45^ have found that exposure to 400 mJ/cm^2^ of this broad-spectrum UV provides the equivalent of 2 MEDs to mice, resulting in mild burning. In contrast, 3 exposures to 20mJ/cm^2^ provides a sub-erythemal dose (0.2 MEDs) that causes photoadaptation, including skin darkening and photoprotection^45,69^ in C57Bl/6 mice. Alternatively for UVA and UVA bandwidth specific experiments, mice were exposed to a UVA or UVB narrowband lamp (DermaHealer® compact; UAB Favoriteplus; Lithuania). In all cases, lamp output was calibrated using a UV505 UV-AB light meter (Extech Instruments).

### Mosquito biting and virus infection

To ensure mosquitoes bit a defined area of skin (upper/dorsal side of the left foot), anesthetized mice were placed onto a mosquito cage containing *Ae. aegypti* mosquitoes and the rest of the body was protected^6^. Biting was monitored and mice were removed once two to five mosquitos were seen to be engorged from feeding. Where required, bitten skin was immediately injected with virus in a 0.5 to 1 μl volume into the skin using a Hamilton Syringe (Hamilton) using either; 1×10^4^ PFU SFV4, 1×10^3^ SFV6-mCHERRY or 1×10^4^ PFU ZIKV. For survival curves, mice were monitored closely and culled when they reached clinically defined endpoints of disease.

### Vitamin D and steroid treatments

Vitamin D (25(OH) D) (Sigma-Aldrich, H4014-1MG) was reconstituted to 1mg/ml in 100% ethanol and then further diluted to 5μg/ml in mineral oil. Mice were injected s.c. with 5ng of vitamin D at UV-exposed site 1 h post-exposure or resting skin. A steroid cream, clobetasol propionate corticosteroid (Dermovate Ointment, GSK) was applied topically to UV-exposed skin or resting skin twice daily for 5 days, with the first treatment applied 1 h post-exposure.

### Oedema quantification

Mice were injected s.c. in the loose dorsal flank/inter-scapular skin with 200μl of 1% Evan’s Blue dye (Sigma-Aldrich) at a set time point post-UV exposure, and 1 h post injection, once blood/systemic levels of dye had been established, were exposed to mosquito bites if required. Skin and blood samples were collected 3 h post-dye injection. Quantity of Evan’s blue in skin was quantified using colorimetric measurement at 620 nm using the Cytation 5 Imaging Reader (Biotek) and normalised to levels in serum.

### Analysis of skin temperature and mosquito behaviours

A Testo 830-T1 Infrared Thermometer (Testo) was used to measure the temperature of UV-exposed or unexposed mouse skin at set time points post-UV exposure. ‘Landing time’ was measured as the time period between when a mouse was set down on top of the mosquito cage with the skin being exposed to potential mosquito bites to when a mosquito landed on the skin. ‘Probing time’ was measured as the time period between when a mosquito began probing the mouse skin looking for a blood meal to when the mosquito abdomen was fully engorged with blood and disengaged from the mouse.

### Gene expression analysis and histology

RNA was extracted using PureLink RNA Mini (Invitrogen) columns and reverse transcribed to cDNA using the High Capacity RNA-to-cDNA kit (Applied Biosystems). Viral RNA and host gene transcripts were quantified by qRT-PCR and infectious virus by end-point titration, as described previously ^70^. qPCR primers for SFV amplified a section of E1^6^, while primers for ZIKV amplified a section of the env gene. For SFV and ZIKV, qPCR assays measured the sum value of both genome and sub-genomic RNA. For histology, skin was formalin fixed and paraffin embedded (FFPE), sectioned and stained with hematoxylin and eosin. Slides were scanned at 20X magnification using the ZEISS AxioScan Z.1 Slide Scanner (ZEISS). Scanned images were analysed using the open-source software QuPath v0.3.2 (ref 290 from thesis). Skin thickness measurements were taken from 10 areas per condition.

### Processing of skin for FACS, MACS, *in vitro* culture and cell proliferation assays

For FACS, skin tissue samples were enzymatically digested with collagenase, dispase and DNAse, stained with antibodies and a viability dye^6^. Cells from digested skin were separated into CD45+ and CD45-fractions via positive selection using CD45 MicroBeads (Miltenyi Biotec). For *in vitro* culture of murine skin cells, cells from digested skin were cultured in a 96-well plate, pre-coated with 0.2% Gelatin Solution (Sciencell) and, where required, infected *ex vivo* with SFV4 at an MOI of 2. To measure cell proliferation, the Click-iT Plus EdU Alexa Fluor 488 Flow Cytometry Assay Kit (Invitrogen) was used. Briefly, cells from digested skin were incubated with EdU (5’-ethynyl-2’-deoxyuridine), a thymidine nucleoside analogue, for 3 h at 37°C to label proliferating cells. A fluorophore was then attached to EdU using Click-iT chemistry for EdU quantification via flow cytometry. All flow cytometry analysis was performed on a CytoFLEX LX (Beckman Coulter Life Sciences). Monocytes were CD45^+ve^ Ly6G^-ve^ CD11b^+ve^ Ly6C^+ve^; neutrophils were CD45^+ve^ Ly6C^-ve^ CD11b^+ve^ Ly6G^hi^; macrophages were CD45^+ve^ Ly6G^-ve^ CD11b^+ve^ MerTK^+ve^; dendritic cells were CD45^+ve^ Ly6C^-ve^ CD11c^+ve^ MHCII^+ve^; endothelial cells were CD45-^ve^CD326^-ve^Vimentin^-ve^CD31^+ve^), fibroblasts were CD45-^ve^CD326^-ve^Vimentin^+ve^ CD31^-ve^ and epithelial cells were CD45-^ve^ CD326^+ve^ Vimentin^-ve^ CD31^-ve^.

### Statistical analysis

Data were analyzed using Prism Version 9 software. Quantities of virus RNA and infectious titres from virus-infected mice were typically not normally distributed and were accordingly analysed using the non-parametric based tests Mann-Whitney or Kruskal-Wallis test with Dunn’s multiple comparison test where appropriate, unless otherwise stated in figure legends. All such column plots show the median value +/- interquartile range. Where data were normally distributed, data were analysed using ANOVA with Holm-Sidak’s multiple comparison test, or unpaired t-test with Welch’s correction, and plotted with mean value. Survival curves were analysed using the logrank (Mantel Cox) test. All plots have statistical significance indicated; *p<0.05, **p<0.01, ***p<0.001, ****p<0.0001, ns=not significant, or p value stated. Occasional samples were removed based on two strictly applied criteria; if experimental administration of substance (e.g. virus) to skin accidentally ruptured a blood vessel, as previously established^6^; or for qPCR data if 18S expression deviated by more than 4-fold from group median.

### Key resources table

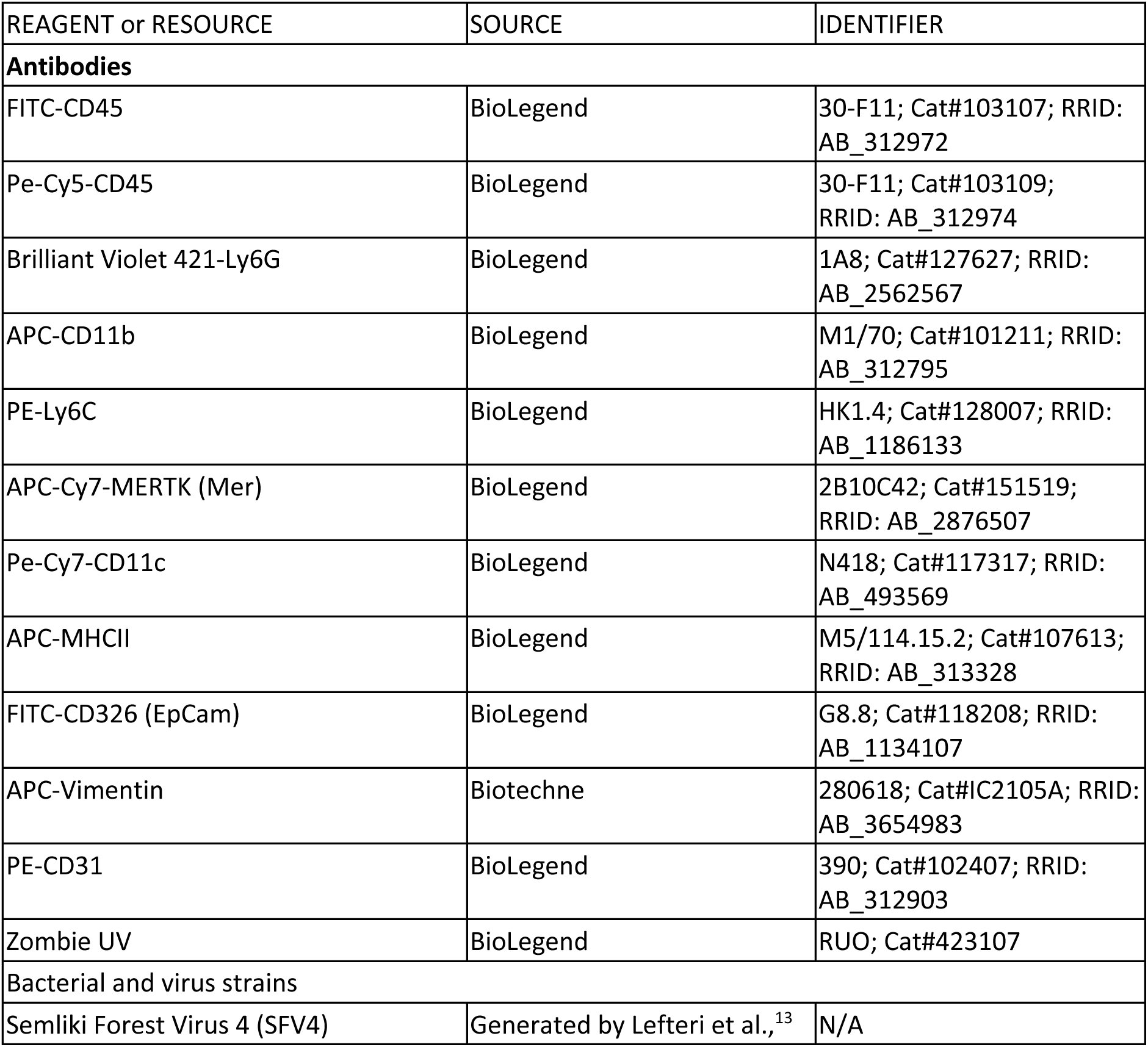
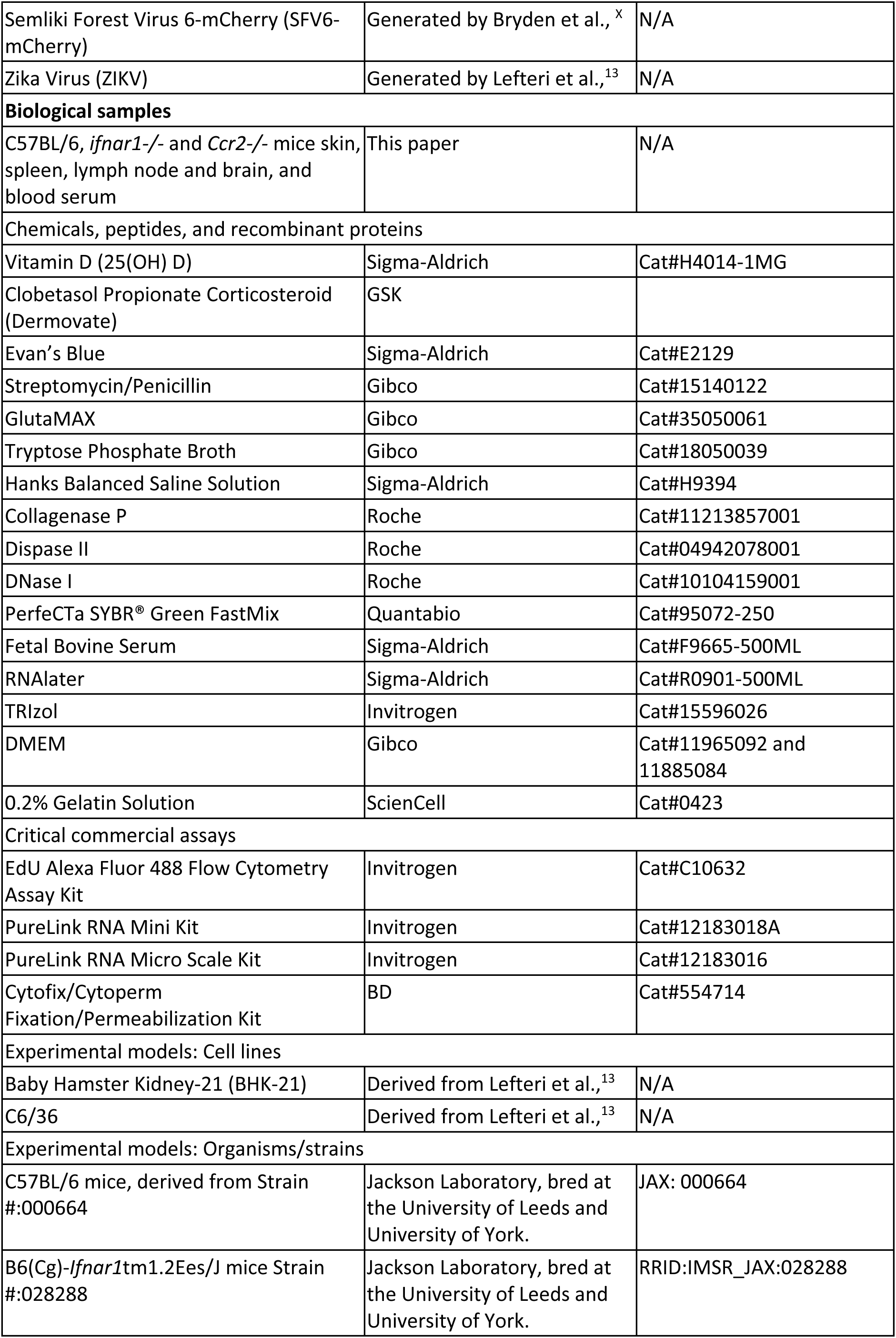
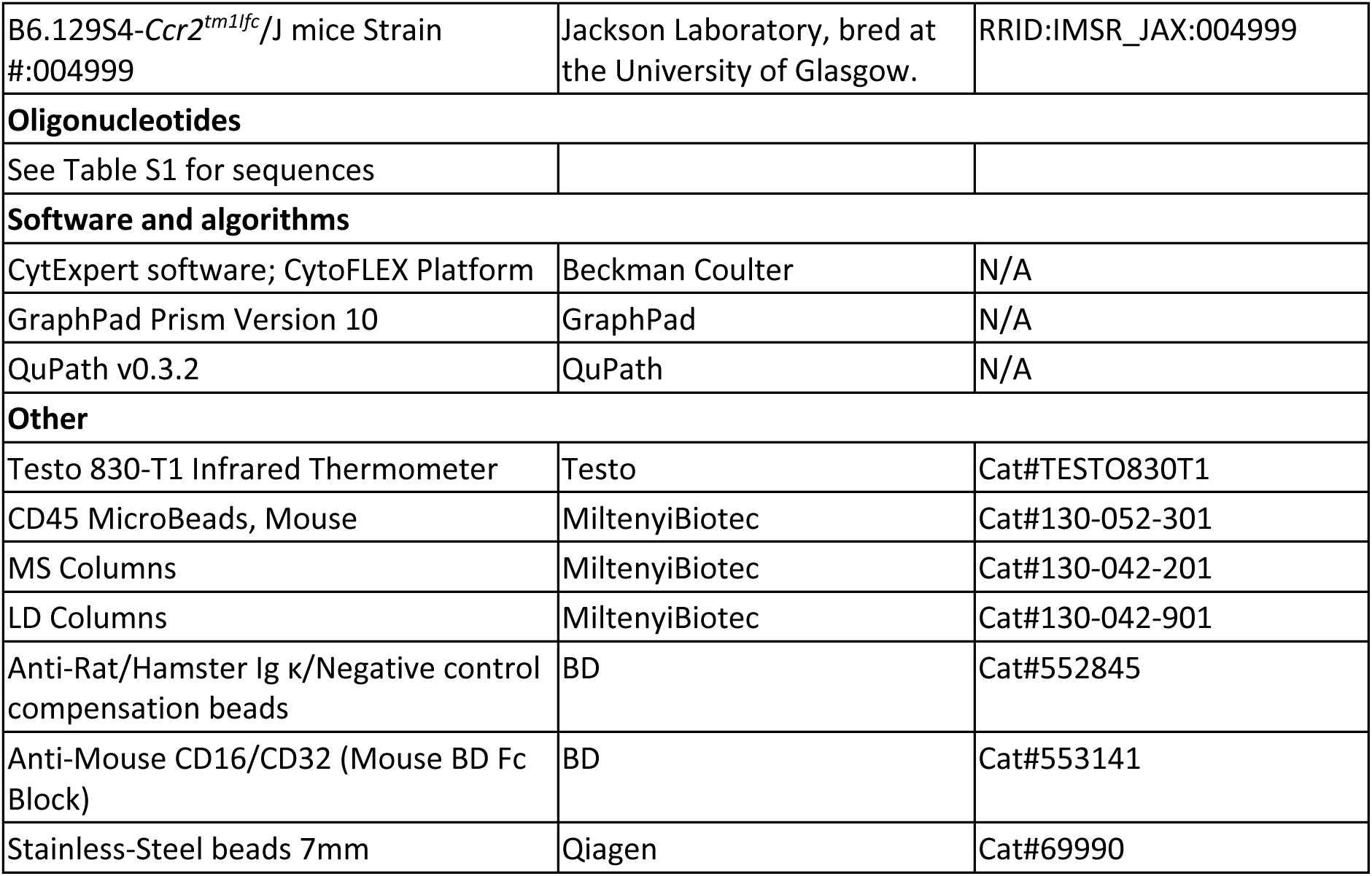

