## Supplementary figures and images for "Ultraviolet exposure conditions skin to enhance arbovirus infection and mosquito probing"

### Figure S1

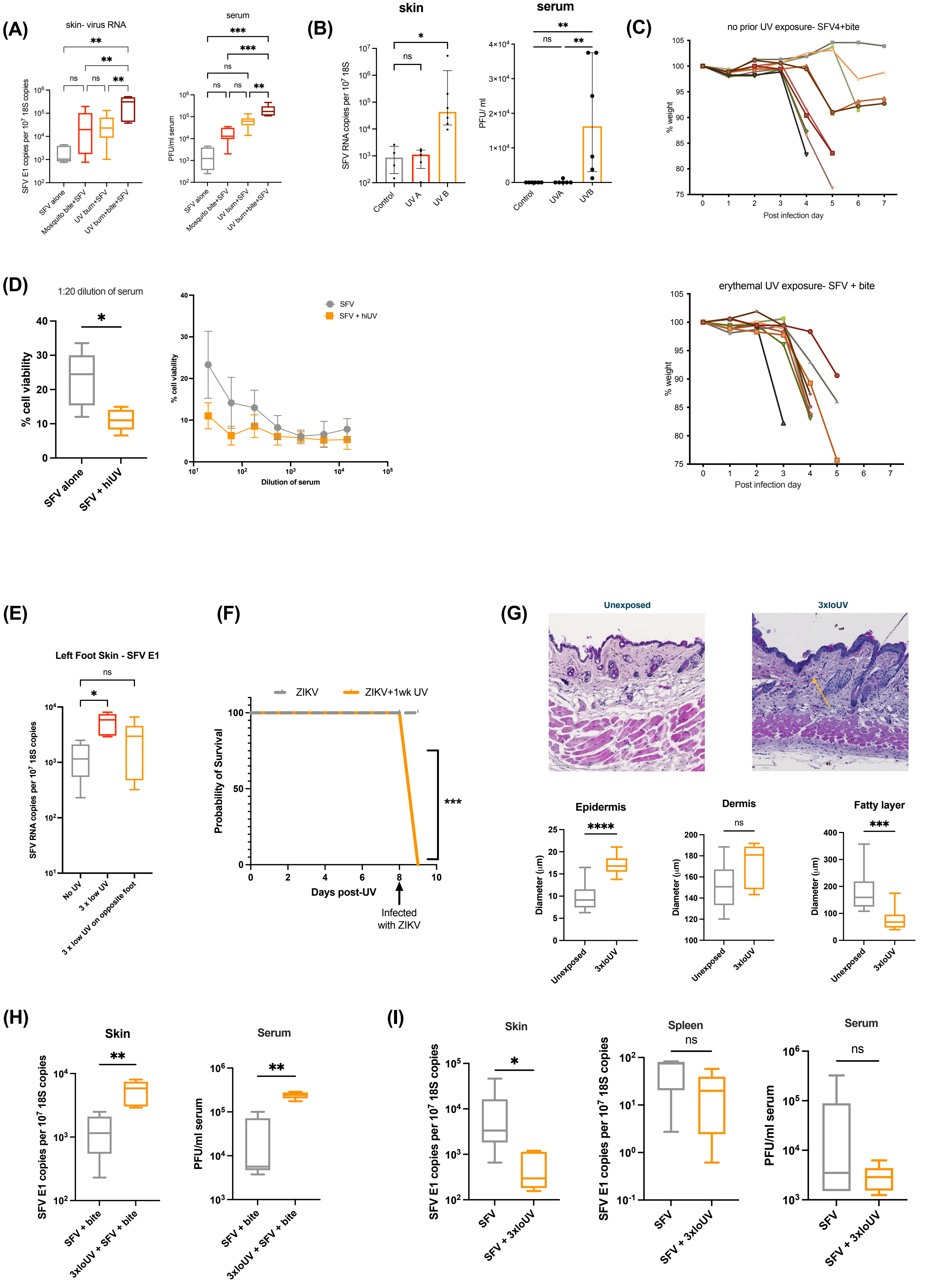

### Figure S2

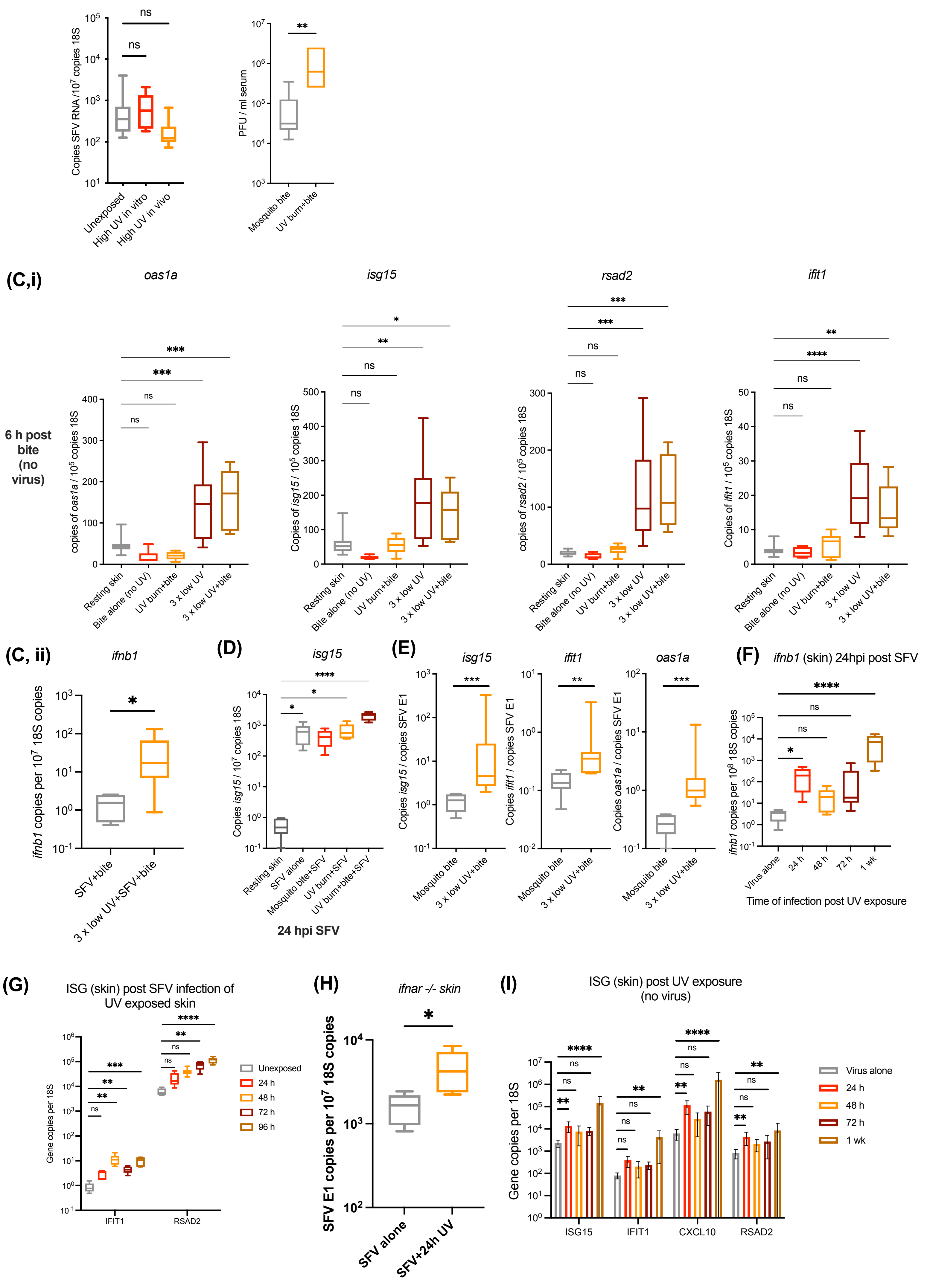

### Figure S3

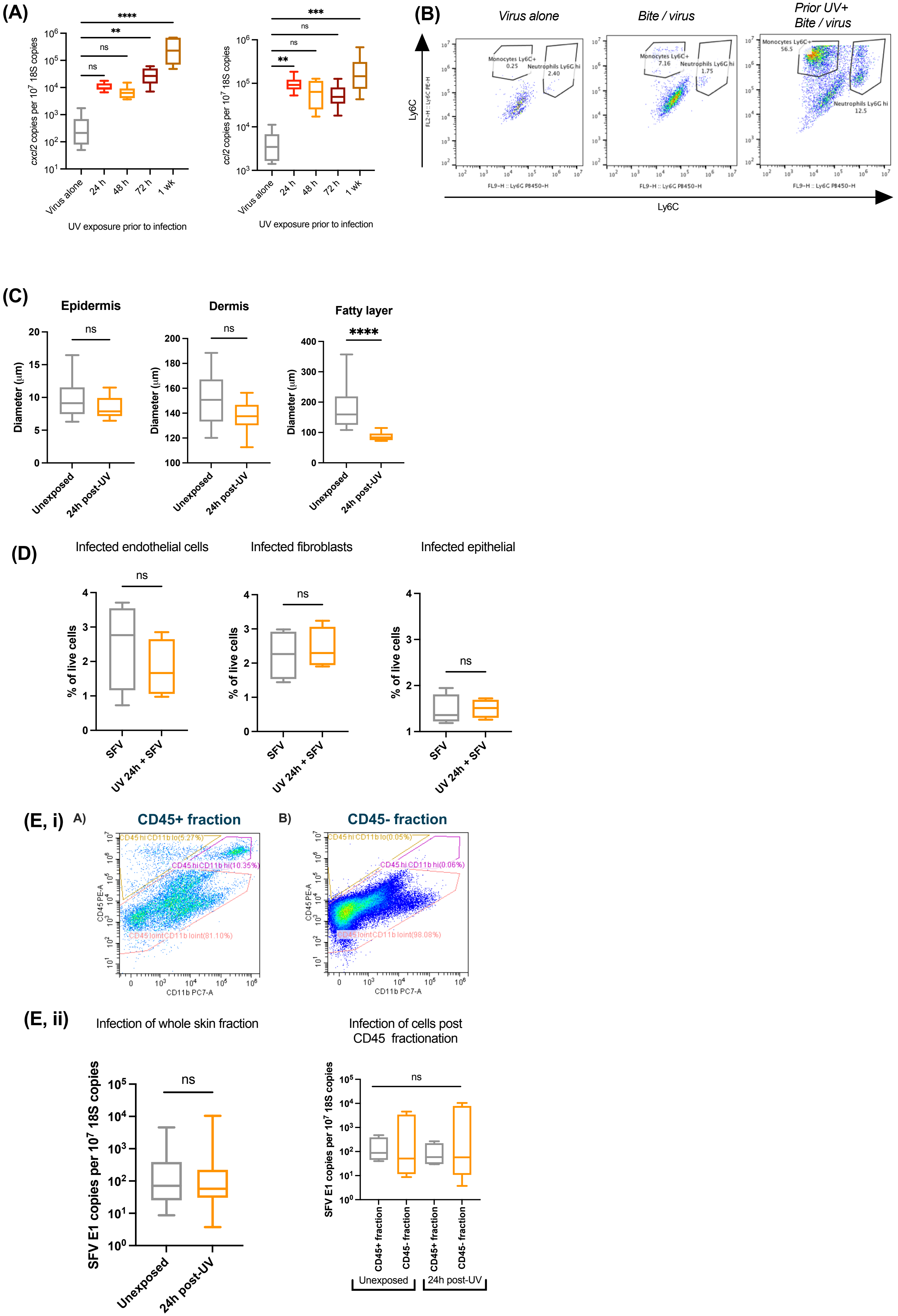

### Figure S4

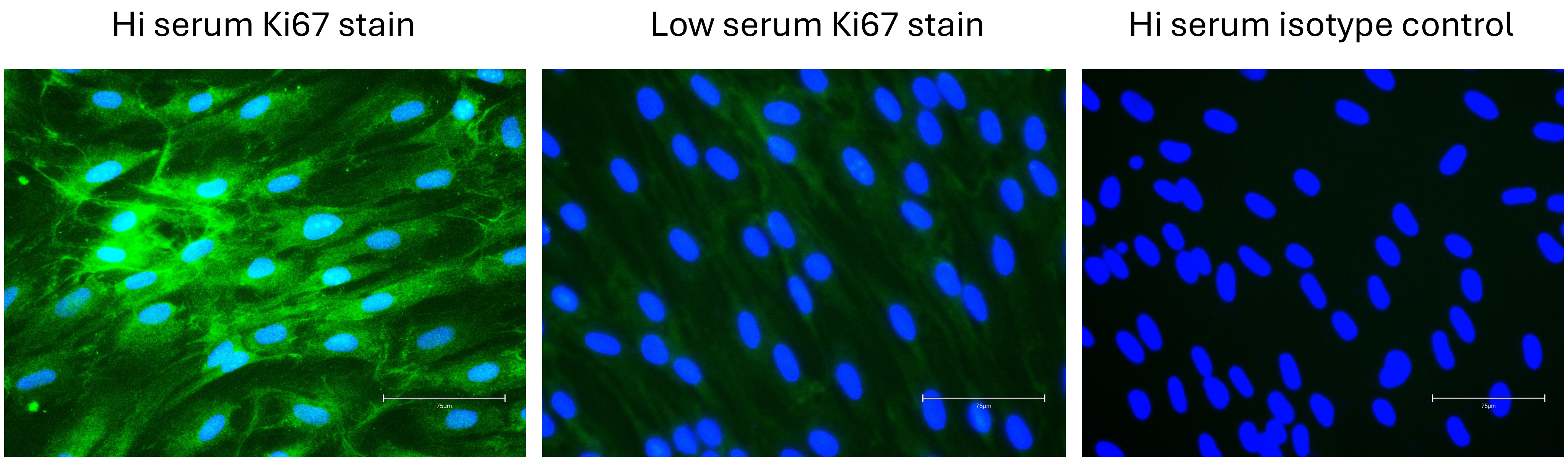

### Figure S5

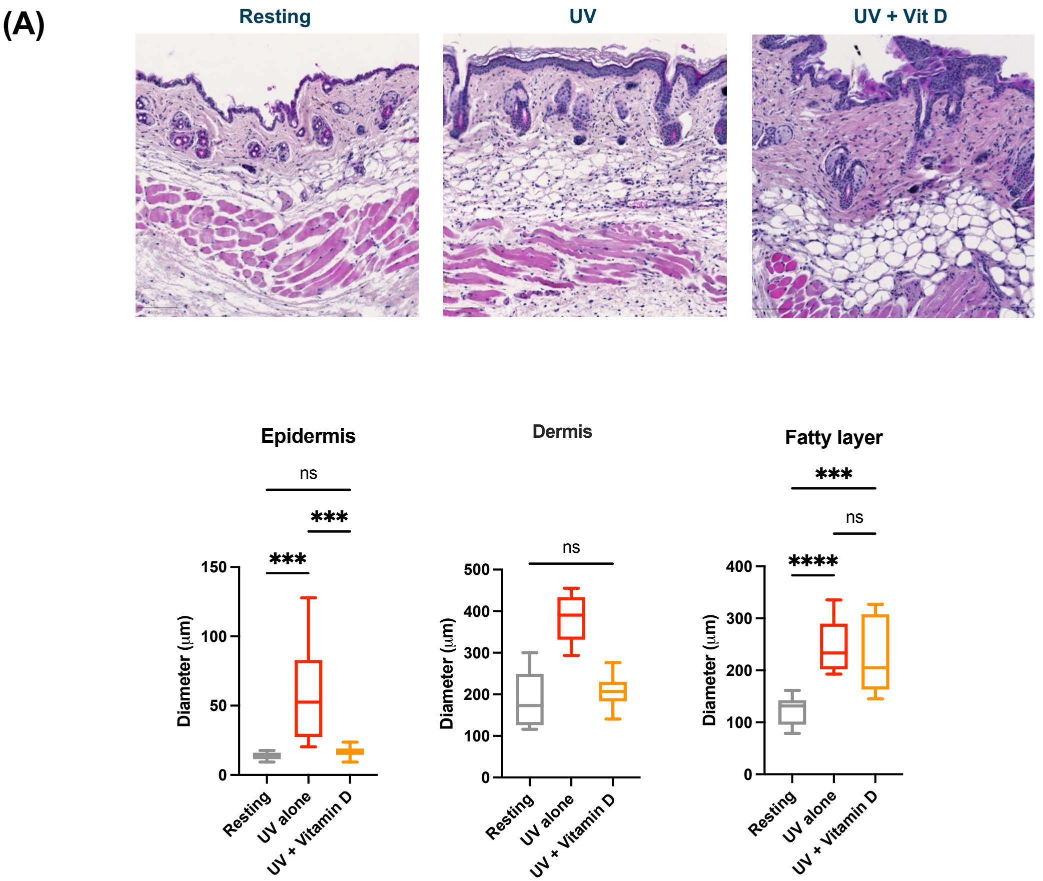

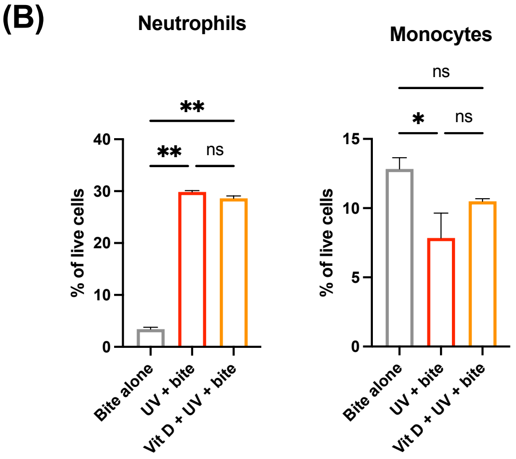

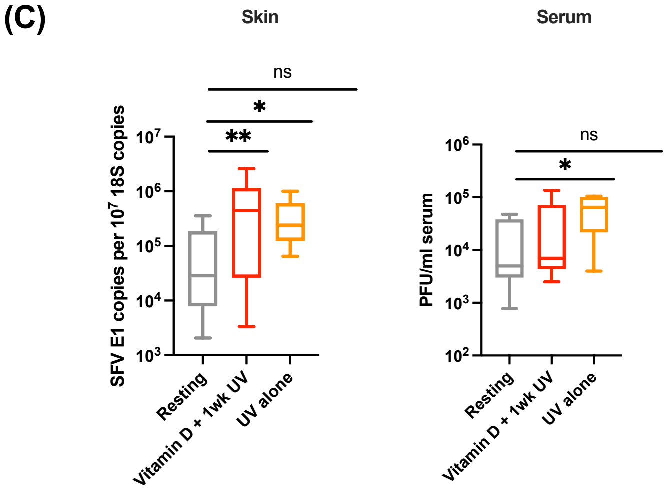

### Figure S6

#### Figure S6

(A)

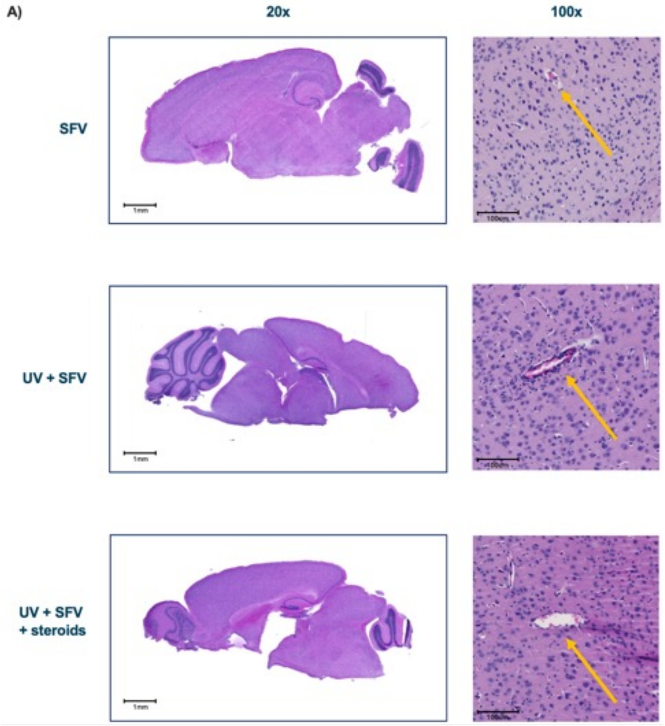


(B)

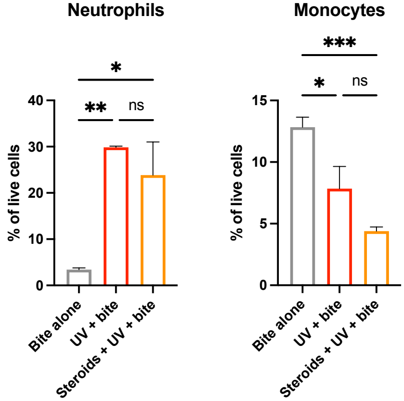


(C)

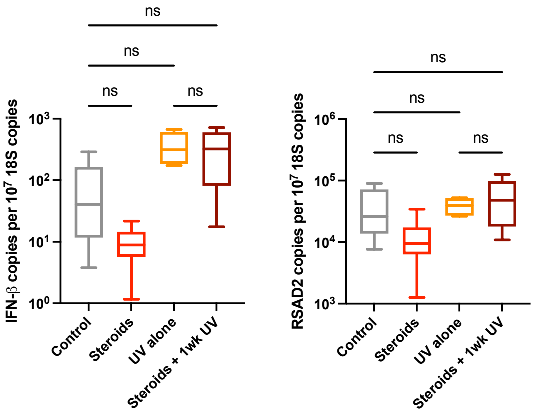
