## Supplementary Figure legend text for "Ultraviolet exposure conditions skin to enhance arbovirus infection and mosquito probing"

### Supplementary Figures

#### Figure S1. Related to Figure 1.

(A) Additional representative experiments in which mice were exposed to erythemal UV and infected with SFV at mosquito bites 24 h later. Viral RNA was quantified by qPCR and serum infectious titres by plaque assay at 24 hpi.
(B) Mice were exposed to 400 mJ/cm² UVA or UVB using wavelength-specific lamps and infected with SFV 1 week later. Viral RNA was quantified by qPCR and serum infectious titres by plaque assay at 24 hpi.
(C) Mouse weight was assessed daily for 7 days after infection. Missing data points indicate mice that were culled after reaching defined humane endpoints.
(D) Mice were infected with SFV at unexposed or UV-exposed skin. Serum collected 1 week post-infection was UV inactivated and incubated with SFV stocks to assess neutralising capacity. Infectious titres were quantified by plaque assay on BHK cells.
(E) Mice were exposed to repeated non-erythemal UV every 48 h for 6 days and infected with SFV 24 h after the final exposure, with mosquito biting, either at the UV-exposed site or at an unexposed site on the contralateral foot. Viral RNA was quantified by qPCR at 24 hpi.
(F) Survival of ZIKV-infected mice following infection of UV-exposed or non-UV-exposed mosquito-bitten skin 1 week post-UV.
(G) Histological assessment and quantification of skin compartment thickness in mouse skin exposed to repeated non-erythemal UV every 48 h for 6 days and sampled 24 h after the final exposure.
(H,I) Additional representative experiments in which mice were exposed to repeated non-erythemal UV every 48 h for 6 days and infected 24 h after the final exposure at the UV-exposed site, either with (H) or without (I) prior mosquito biting. Viral RNA was quantified by qPCR at 24 hpi and serum infectious titres by plaque assay.
Note: In panel D, double-check whether "serum was inactivated" means heat-inactivated, UV-inactivated, or something else. "UV-inactivated serum" reads oddly unless that was truly the method.

#### Figure S2. Related to Figure 2. qPCR analysis of *ifnb1* and ISGs following UV exposure, mosquito biting and SFV infection.

(A) Mouse skin explants were exposed to erythemal UV either in vivo or ex vivo in tissue culture, or left unexposed, then infected ex vivo with $1 \times 10^5$ PFU SFV. Viral RNA was quantified by qPCR at 24 hpi (n = 8).
(B) NSG immunodeficient mice were exposed to erythemal UV and infected with SFV at mosquito bites 24 h later. Viral RNA was quantified by qPCR at 24 hpi (n = 6).
(C) Mice were exposed to a single erythemal UV dose, repeated non-erythemal UV every 48 h for 6 days, or left unexposed. Twenty-four hours after the final UV exposure, mice were either left untreated or exposed to mosquito bites, and gene expression was assessed by qPCR 6 h later (i), or infected with SFV and *ifnb1* expression quantified at 24 hpi (ii) (n = 6).
(D) Mice were exposed to a single erythemal UV dose or left unexposed. Twenty-four hours later, mice were infected with SFV alone or alongside mosquito bites, and *isg15* expression was quantified by qPCR at 24 hpi (n = 6).
(E) Mice were exposed to repeated non-erythemal UV every 48 h for 6 days or left unexposed. Twenty-four hours after the final UV exposure, mice were bitten by mosquitoes, and gene expression was assessed by qPCR 6 h later (n = 8).

(F) Mice were exposed to a single erythemal UV dose or left unexposed. Mice were infected with SFV at the indicated times post-UV exposure, and *ifnb1* transcripts were quantified by qPCR at the indicated time points post-infection (n = 6).
(G) Mice were exposed to a single erythemal UV dose or left unexposed. At the indicated times post-UV exposure, mice were infected with SFV at mosquito bites and gene transcripts were quantified by qPCR at 24 hpi (n = 6).
(H) *Ifnar1*-null mice were exposed to erythemal UV and infected with SFV at mosquito bites 24 h later. SFV RNA was quantified by qPCR at 24 hpi (n = 5).
(I) Mice were exposed to a single erythemal UV dose or left unexposed. At the indicated times post-UV exposure, gene transcripts were quantified by qPCR (n = 6).
Statistical significance: *p < 0.05, **p < 0.01, ***p < 0.001, ****p < 0.0001, ns = not significant; Mann-Whitney test or Kruskal-Wallis test with Dunn's post-test for comparisons of three or more groups.

**Figure S3. Related to Figure 2.**
(A) Mice were exposed to a single erythemal UV dose or left unexposed. At the indicated times post-UV exposure, mice were infected with SFV at mosquito bites, and chemokine transcripts were quantified by qPCR at 24 hpi (n = 6).
(B) Representative flow-cytometry gating for skin monocytes ($CD45^+CD11b^+Ly6C^+Ly6G^-$) and neutrophils ($CD45^+CD11b^+Ly6C^-Ly6G^+$) at 16 hpi with SFV4 in erythemal UV-exposed, mosquito-bitten skin.
(C) Skin compartment thickness measured from H&E-stained sections of erythemal UV-exposed skin at 24 hpi.
(D) Frequency of $mCherry^+$ cells at 24 hpi following infection with SFV6-mCherry in erythemal UV-exposed, mosquito-bitten skin (n = 4).
(E) Representative flow-cytometry gating showing enrichment and depletion of the $CD45^+$ fraction following magnetic bead-based separation (i). Viral RNA was quantified by qPCR in cells infected in vitro with $1 \times 10^4$ PFU SFV before or after CD45 fractionation, following isolation from mice left unexposed or exposed to erythemal UV in vivo 24 h before isolation (ii) (n = 8).
Statistical significance: *p < 0.05, **p < 0.01, ***p < 0.001, ****p < 0.0001, ns = not significant; Mann-Whitney test or Kruskal-Wallis test with Dunn's post-test for comparisons of three or more groups.

**Figure S4. Low-serum culture conditions reduce fibroblast Ki67 expression.**
Primary human dermal fibroblasts (HDFs) were cultured in high-serum (10% FBS) or low-serum (0.5% FBS) conditions for 24 h, stained for Ki67 (green) with DAPI nuclear counterstain (blue), and imaged by EVOS microscopy. Representative images show reduced Ki67-positive cells under low-serum conditions (n = 8). Scale bars, 75 μm.

**Figure S5. Vitamin D treatment of UV burn.**
(A) Histological assessment showing partial reversal of UV-induced epidermal thickening by vitamin D treatment. Thickness of each skin compartment was measured.
(B) Flow-cytometric quantification of neutrophil and monocyte frequencies following vitamin D treatment (n = 3). Monocytes were defined as $CD45^+CD11b^+Ly6C^+Ly6G^-$ and neutrophils as $CD45^+CD11b^+Ly6C^-Ly6G^+$.

(C) Mice were exposed to erythemal UV or left unexposed, then treated or not treated with vitamin D (n = 6). At 1 week post-UV exposure, mice were infected with SFV, and viral RNA and serum infectious titres were quantified at 24 hpi by qPCR and plaque assay, respectively. Statistical significance: *$p < 0.05$, **$p < 0.01$, ***$p < 0.001$, ****$p < 0.0001$, ns = not significant; Mann-Whitney test or Kruskal-Wallis test with Dunn's post-test for comparisons of three or more groups.

**Figure S6. Related to Figure 6. Steroid treatment of UV burn.**
(A) Brain histology showing perivascular cellular infiltrates in the cortex after UV exposure, reduced by steroid treatment.
(B) Skin leukocyte frequencies in mice exposed to erythemal UV and then left untreated or treated with topical steroid for 6 days, assessed by flow cytometry (n = 3).
(C) Skin *Ifnb1* and *Rsad2* expression quantified by qPCR 1 week after erythemal UV exposure, with or without steroid treatment (n = 6).
Statistical significance: *$p < 0.05$, **$p < 0.01$, ***$p < 0.001$, ****$p < 0.0001$, ns = not significant; Mann-Whitney test or Kruskal-Wallis test with Dunn's post-test for comparisons of three or more groups.
